# Genetic divergence outpaces phenotypic divergence among threespine stickleback populations in old freshwater habitats

**DOI:** 10.1101/618017

**Authors:** Mark C. Currey, Susan L. Bassham, William A. Cresko

**Affiliations:** Institute of Ecology and Evolution, University of Oregon, Eugene, OR 97403-1254

**Keywords:** Gasterosteus aculeatus, RAD-seq, principle component analysis, population structure

## Abstract

Species such as threespine stickleback fish that are distributed across landscapes with divergent selective environments and that have diversified on different time scales can be valuable for understanding evolutionary processes. Here we synthesize high resolution genotypic and phenotypic data to explore a largely unstudied distribution of threespine stickleback populations living in marine and freshwater habitats along coastal and inland regions of northwestern Oregon. Because many inland aquatic habitats of Oregon were not glaciated during the last ice age, we hypothesized that some extant Oregon lake and river stickleback are descended from freshwater populations that were established long before the well-studied, postglacial freshwater populations of Alaska. Here we characterize the major phenotypic and genetic axes of differentiation in Oregon stickleback, and compare these patterns to their Alaska counterparts currently inhabiting regions that were covered by ice during the last glacial maximum. Phenotypic variation in Oregon stickleback is predictably partitioned between oceanic and freshwater habitats. However, we also found that genetic divergence in Oregon ecotypes is much greater than divergence among studied stickleback populations in Alaska. Additionally, we report a surprising phenotypic and genetic affinity between oceanic stickleback with freshwater populations that live far inland in two Oregon river basins.

## INTRODUCTION

Studies that pair the partitioning of phenotypic and genetic variation within and among populations have long been used to infer ecological and evolutionary processes from patterns observed in nature (Endler, 1986; Storz, 2002; Nosil *et al.,* 2008; Nosil & Schluter, 2011). Integration of these data types can be potent for biological inference, particularly when data are collected from the same set of individuals in populations that occupy diverse habitats and especially where parallel phenotypic evolution manifests across geographic locations and/or geological timescales (Endler, 1977; Endler, 1986; Schluter, 2000). Recent studies that unite population genomic and phenotypic data from hundreds or thousands of individuals provide novel insights into the direction and pace of genetic and phenotypic evolution (Nosil *et al.,* 2002; Nadeau *et al.,* 2012; Soria-Carrasco *et al.,* 2014; Ferrero-Serrano & Assmann, 2019). Such combined data sets are ushering in a new age of natural history that includes the mapping of genetic diversity to geographic and phenotypic variation in such fields as conservation genetics (Funk *et al.,* 2012; Casillas & Barbadilla, 2017), population genomics (Andrews *et al.,* 2016), landscape genetics (Dionne *et al.,* 2008; McCairns & Bernatchez, 2008; Gaggiotti *et al.,* 2009), and genome-wide association studies (Pallares *et al.,* 2014; Gienapp *et al.,* 2017; Saltz *et al.,* 2017).

A powerful way to implement this new synthesis is to perform population genomic analyses in an organism for which extensive data on the ecology, evolution, and biogeography have already been gathered. The threespine stickleback fish *(Gasterosteus aculeatus)* is one such system. This small fish is distributed holarctically, and is abundantly present in oceanic, estuarine, and freshwater habitats, encompassing populations with phenotypically diverse life history forms in nearly all parts of the coastline in the northern half of the Northern Hemisphere (Bell & Foster, 1994). Consequently, the species has for decades been a rewarding focus of studies in behavior, ecology, physiology, ecotoxicology and evolution (e.g., Tinbergen, 1950; Foster, 1995; Bernhardt *et al.,* 2006; Hohenlohe *et al.,* 2010; Jones *et al.,* 2012a; Reimchen *et al.,* 2013; Spence *et al.,* 2013; Furin *et al.,* 2015; Teigen *et al.,* 2015; Divino *et al.,* 2016; Greenwood *et al.,* 2016; Bassham *et al.,* 2018; Hani *et al.,* 2018). The natural history of the species is therefore one of the best documented of any vertebrate, and the ecological and evolutionary processes that create and maintain its diversity across a variety of biological levels are increasingly well understood (e.g., Colosimo *et al.,* 2004; Cresko *et al.,* 2004; Hohenlohe *et al.,* 2010; Schluter *et al.,* 2010; Jones *et al.,* 2012a; Bassham *et al.,* 2018).

An exceptional attribute of the stickleback species is its repeated ecological transition across profoundly different environments. Anadromous stickleback populations have episodically colonized and adapted to freshwater habitats (Schluter & Conte, 2009; Bell & Aguirre, 2013), likely throughout time since the origin of the species (Bell & Foster, 1994; Bell, 2009; Bell *et al.,* 2009) and continuing to the present day as new habitats are created (Hagen & Gilbertson, 1972; Klepaker, 1993; Bell, 2001; Bell *et al.,* 2004; Bell & Aguirre, 2013; Lescak *et al.,* 2015). This long history has given rise to a large set of replicate natural evolutionary experiments. Phenotypic diversification in these fish spans morphological, behavioral, physiological, and life history traits. Among these, the best documented are structural phenomena, including change in defensive armor, such as loss of lateral plates and reduction or loss of the pelvic structure, differences in body shape related to foraging or predation avoidance, and changes in craniofacial morphology linked to alternative foraging ecologies (Hagen & Gilbertson, 1972; Bell & Foster, 1994; Walker, 1997; Kimmel *et al.,* 2012; Wund *et al.,* 2015).

Surprisingly, at least some of the parallel phenotypic divergence observed in many of these populations appears to have a parallel genetic basis (Cresko *et al.,* 2004; Colosimo *et al.,* 2005; Hohenlohe *et al.,* 2010; Deagle *et al.,* 2012; Jones *et al.,* 2012b; Bassham *et al.,* 2018). Recent work shows that not only can these phenotypic and genetic changes happen in the thousands of years since the end of the last glacial maximum (Bell & Orti, 1994; Rundle *et al.,* 2000; Cresko *et al.,* 2004; Hohenlohe *et al.,* 2010; Hendry *et al.,* 2011; Reimchen *et al.,* 2013), but that significant changes can occur in freshwater populations founded even just decades ago (Bell *et al.,* 2004; Kitano *et al.,* 2008; Gelmond *et al.,* 2009; Lucek *et al.,* 2014; Lescak *et al.,* 2015; Bassham *et al.,* 2018).

Despite the rapid advances in stickleback population genomics, the majority of work on stickleback has been performed in populations spread throughout the higher latitudes of the northern hemisphere in relatively young freshwater aquatic habitats that arose since the last glacial maximum. Much less work has been performed in older habitats at lower latitudes that were not subject to the most recent glacial events (but see Hagen & Gilbertson, 1972; Bell & Richkind, 1981; Baumgartner & Bell, 1984; Sanchez-Gonzales *et al.,* 2001; Morris *et al.,* 2018). How does the phenotypic and genetic diversity of stickleback in these older habitats compare to well-studied stickleback populations from younger habitats? And, is there a roughly linear relationship in increasing phenotypic and genetic divergence over time? To address these questions, we gathered extensive phenotypic and population genomic data from stickleback populations that inhabit the aquatic habitats of Oregon, and for which little is known (Rutter, 1896; but see Catchen *et al.,* 2013).

The unique geography and geologic history of Oregon has created an array of aquatic habitats of varying ages and degrees of connectivity with the ocean, characterized by young freshwater habitats along the Oregon coast and much older inland habitats. There are 22 major estuaries along the Oregon coast, many of which were formed when the mouths of their associated rivers were drowned by rising sea level during the last glacial maximum (Allen & Baldwin, 1944). The coast line is also dotted with many freshwater lakes, some of which were formed during the Holocene (0.1-7 ka) by growing sand dunes that blocked run-off (Peterson *et al.,* 2007). Often these coastal ponds outflow to the sea via short channels. In contrast, Oregon’s inland stickleback habitats can be hundreds of miles from the sea, along the Willamette, Deschutes, and Umpqua River basins, which have existed in roughly their present forms for millions of years (Booth *et al.,* 2003). Previous research on stickleback divergence suggests that much of the genetic variation important for initial adaptation to fresh water is carried by - but not expressed in - marine populations and predates the last glacial maximum (Schluter & Conte, 2009; Hohenlohe *et al.,* 2010; Terekhanova *et al.,* 2014; Bassham *et al.,* 2018; Nelson & Cresko, 2018). We therefore expect to find broadly similar patterns of phenotypic change from the marine form in both young and old Oregon freshwater populations and younger Alaska freshwater populations, but the existence of potentially more isolated and much older inland freshwater populations in Oregon predicts that levels of genetic differentiation could be more extreme than those found in Alaska.

Here we integrate morphometrics of defensive, trophic, and body size traits with RAD-seq population genomic data collected from thousands of individual stickleback disbursed over many populations. To understand how phenotypic and genetic variation is partitioned across extensive landscapes that were not under ice during the last glacial maximum we compare populations sampled throughout the northwestern quadrant of Oregon. We then compare patterns of diversification in Oregon to those documented among oceanic and freshwater populations from Alaska. We show that while phenotypic diversification is roughly similar among populations in Oregon and Alaska, we found much greater genetic divergence among some of the freshwater Oregon populations. In contrast to previous studies where the major axis of genetic divergence is between ocean and freshwater populations, we found that the major population genomic split in Oregon is between coastal and inland populations regardless of habitat or life history. We also show that despite connectivity of river systems within the Willamette Basin, we could detect population genetic structuring by distance and altitude, a pattern broadly similar to that seen in other riverine fish, such as observed in Trinidadian guppies and anadromous brook charr. Finally, we document a surprising discovery of stickleback in far inland freshwater habitats that are phenotypically and genetically similar to oceanic types.

## METHODS

### Collection and processing of stickleback samples

Oceanic, coastal freshwater, and inland freshwater populations were collected throughout the northwest region of Oregon. Oceanic and freshwater Alaska populations were collected from Middleton Island, locations in the Matanuska-Sustina valley, and the Kenai peninsula. Some populations from all locations were used in previous studies (Fig. 1, Table 1) (Catchen *et al.,* 2013; Lescak *et al.,* 2015; Bassham *et al.,* 2018). Most stickleback samples were collected using minnow traps as previously described (Cresko *et al.,* 2004; Catchen *et al.,* 2013). A subset were obtained from state and federal agencies, namely Oregon Department of Fish and Wildlife (ODFW) and National Oceanic and Atmospheric Administration (NOAA), which were conducting collections of other fish species in Oregon using various methods of collection. GPS coordinates of collecting locations were obtained using Google Earth. Fish were euthanized and immediately fixed in 95% ETOH. For all samples, the soma of each individual was fixed in 4% PFA overnight at room temperature and then stained with Alizarin Red for morphometric analysis (Cresko *et al.,* 2004; Lescak *et al.,* 2015). For genetic analyses, tissue samples of both pectoral and caudal fins were collected fresh or after fixing in ETOH and stored at −20° or −80° C or were immediately processed for DNA extraction and subsequent RAD sequencing. All Oregon and most Alaska collections were made between 2007 and 2017, with some of the Alaska samples collected in the mid-1990s. These efforts provided a total of 1419 individuals in 47 populations used in this study. (Fig. 1, Table 1). All research was approved by the Institutional Animal Care and Use Committees (IACUCs) of the University of Alaska Anchorage and the University of Oregon. Fish were collected under Alaska Department of Fish and Game permits SF-2011-153 and SF2014-035 and Oregon Department of Fish and Wildlife scientific taking permits OR2007-3495, 13920, 16933, 17664, 19122, and 20770.

**Figure 1.**
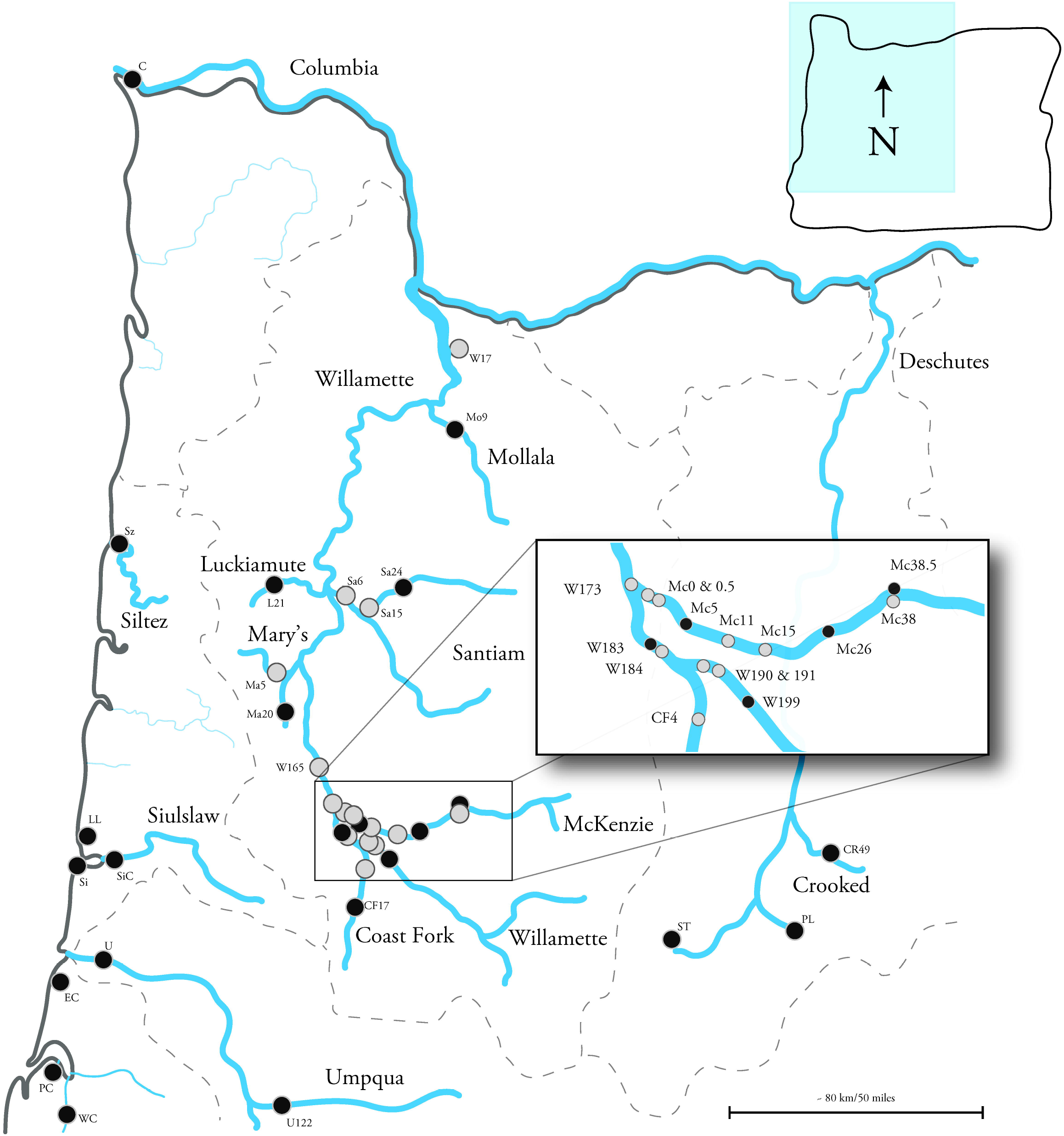
Regional distribution of Oregon populations of threespine stickleback used for phenotypic and genetic analysis. Black dots denote populations used in both analyses, grey dots are populations used in the phenotypic analysis only. Major rivers are shown in blue. Grey dashed lines are major watershed boundaries. Population abbreviations are found in table 1. Numbers associated with population abbreviations denote the number of river miles of each collecting site according to USGS topo maps.

**Table 1.** Collection sites with abbreviation used in figures, major basin or river the site is located in, elevation, distance to ocean, GPS coordinates, collection dates, number of individuals used in genotypic analysis, date sequenced and length of read, and number of individuals from each populations used in the phenotypic analysis.

| Collection Site | Ab | Major Basin/River | Elevation (m/ft) | Distance to Ocean (km/miles) | GPS coordinates | Collection date(s) | Genotyped | Date Sequenced Length | Phenotyped |
| --- | --- | --- | --- | --- | --- | --- | --- | --- | --- |
| Columbia River | C | Columbia | 0/0 | 11/7 | 46°14'36.04"N 123°54'9.11"W | Jul, 2012 | 48 | 2013 – 100 bp | 60 |
| Pony Creek Reservoir | PC | Coos | 29/95 | 23/14 | 43°22'12.03"N 124°15'43.52"W | Mar, 2007 | 70 | (Catchen *et al.* 2013) | 30 |
| Winchester Creek | WC | Coos | 2/8 | 10/6 | 43°16'37.39"N 124°19'8.57"W | 2007 | 22 | (Catchen *et al.* 2013) | 30 |
| Crooked River | CR49 | Crooked R. | 1075/3530 | 589/366 | 44°17'25.57"N 120°50'47.07"W | Aug, 2008 | 30 | (Catchen *et al.* 2013) | 30 |
| Paulina Lake | PL | Deschutes | 1947/6389 | ∞ | 43°42'48.66"N 121°16'21.48"W | Aug, 2008 | 21 | (Catchen *et al.* 2013) | 30 |
| South Twin Lake | ST | Deschutes | 1323/4340 | ∞ | 43°42'38.38"N 121°46'4.53"W | Jun, 2007 | 50 | (Catchen *et al.* 2013) | 30 |
| Jont Creek | L21 | Luckiamute | 68/229 | 370/230 | 44°46'28.32"N 123°19'5.02"W | Oct, 2012 | 48 | 2013 – 100 bp | 30 |
| Dunawi Creek | Ma5 | Mary's | 73/239 | 383/238 | 44°32'55.04"N 123°18'9.66"W | Aug, 2013 |  |  | 30 |
| Finley Grey Creek Swamp | Ma20 | Mary's | 95/312 | 407/253 | 44°23'44.09"N 123°20'35.42"W | Aug, 2013 | 48 | 2013 – 100 bp | 29 |
| Bear Paw Lake | BP | Mat-Su Ak | 85/281 | 20/13 | 61°36'53.79"N 149°45'23.32"W | June, 2004 | 47 | 2018 – 150 bp | 20 |
| Mud Lake | Md28 | Mat-Su Ak | 30/97 | 82/51 | 61°56'8.27"N 150°58'44.01"W | 1997 |  |  | 20 |
| Meadow Creek | Me8 | Mat-Su Ak | 426/142 | 13/8 | 61°33'39.87"N 149°50'0.33"W | July, 1996 |  |  | 20 |
| Rabbit Slough | Ra1 | Mat-Su Ak | 18/5 | 8/5 | 61°32'10.85"N 149°15'30.01"W | 2014 | 48 | 2017 – 150 bp | 20 |
| Confluence Slough | Mc0 | McKenzie | 113/372 | 445/276 | 44°7'32.21"N 123° 6'22.13"W | Jun, 2013 |  |  | 19 |
| Confluence Island | Mc0.5 | McKenzie | 113/372 | 446/277 | 44°7'22.07"N 123° 6'8.59"W | Jun, 2013 |  |  | 15 |
| Wildish | Mc5 | McKenzie | 128/421 | 452/281 | 44°7'0.53"N 123° 4'17.23"W | Jun, 2013 |  |  | 30 |
| Riverbend | Mc11 | McKenzie | 132/432 | 461/287 | 44°4'40.97"N 123° 1'34.92"W | 2010 - 2013 | 140 | (Catchen *et al.* 2013) | 41 |
| Rainbow Water | Mc15 | McKenzie | 144/475 | 468/291 | 44°3'42.63"N 122°58'7.69"W | Aug, 2012 |  |  | 25 |
| McKenzie Oxbow | Mc22 | McKenzie | 160/525 | 480/298 | 44°3'41.29"N 122°51'10.87"W | Oct, 2012 |  |  | 30 |
| Walterville Slough | Mc26 | McKenzie | 184/605 | 486/302 | 44°4'13.64"N 122°47'54.68"W | Aug, 2012 | 24 | 2013 – 100 bp | 50 |
| Leaburg Fish Hatchery | Mc38 | McKenzie | 224/736 | 505/314 | 44°8'3.84"N 122°36'31.86"W | Oct, 2012 |  |  | 50 |
| Leaburg Lake | Mc38.5 | McKenzie | 227/746 | 507/315 | 44°8'12.99"N 122°36'44.99"W | Oct, 2012 | 48 | 2013 – 100 bp | 30 |
| Middleton Island Fresh | MiF | Mid-Is Ak | 0.3/1 | ∞ | 59°27'44.82"N 146°17'53.36"W | May, 2010 | 50 | (Lescak *et al.* 2015) | 20 |
| Middleton Island Oceanic | MiO | Mid-Is Ak | 0/0 | 0/0 | 59°27'39.60"N 146°17'45.60"W | May, 2010 | 32 | (Lescak *et al.* 2015) | 20 |
| Milk Creek | Mo9 | Mollala | 48/159 | 235/146 | 45°14'12.82"N 122°37'54.38"W | Oct, 2013 | 32 | 2013 – 100 bp | 30 |
| Resurrection Bay | Re | Resur. Ak | 0/0 | 0/0 | 60° 6'40.98"N 149°24'24.93"W | 1997 |  |  | 20 |
| Salmon Creek | Re2 | Resur. Ak | 12/38 | 3.2/2 | 60° 8'49.76"N 149°24'46.29"W | 1995 |  |  | 20 |
| I-5 Side Channel | Sa6 | Santiam | 58/189 | 346/215 | 44°44'11.80"N 123° 2'56.67"W | Aug, 2013 |  |  | 23 |
| Green's Bridge | Sa15 | Santiam | 76/248 | 360/224 | 44°42'32.23"N 122°58'12.74"W | Sep, 2013 |  |  | 30 |
| Buell-Miller Slough | Sa24 | Santiam | 120/395 | 375/233 | 44°46'12.44"N 122°50'46.96"W | Aug, 2013 | 21 | 2013 – 100 bp | 21 |
| Millport Slough | Sz | Siletz | 1/3 | 5/3 | 44°53'14.68"N 123°59'46.20"W | Apr, 2010 | 68 | (Catchen *et al.* 2013) | 30 |
| Lily Lake | LL | Siuslaw | 4/14 | 1/0.5 | 44°5'33.69"N 124°06'58.81"W | Aug, 2015 | 24 | 2017 – 150 bp | 30 |
| South Jetty | Si | Siuslaw | 1/3 | 2/1 | 44°0'7.76"N 124° 7'59.26"W | March, 2009 | 86 | (Catchen *et al.* 2013) | 30 |
| Cushman Slough | SiC | Siuslaw | 1.625 | 13/8 | 43°59'22.4"N 124°2'42.94"W | March, 2009 | 95 | (Catchen *et al.* 2013) | 31 |
| Eel Creek | EC | Ten Mile | 5/17 | 8/5 | 43°35'13.06"N 124°11'9.96"W | 2007 - 2017 | 24 | 2017 – 150 bp | 27 |
| Dean Creek | U | Umpqua | 4/13 | 26/16 | 43°41'34.36"N 124° 0'1.66"W | June, 2017 | 12 | 2017 – 150 bp | 30 |
| Page Road | U122 | Umpqua | 133/436 | 196/122 | 43°17'2.70"N 123°19'46.66"W | Sept, 2017 | 46 | 2018 – 150 bp | 30 |
| Mt Pisgah Lily Pond | CF4 | Willamette | 156/518 | 470/292 | 44°0'0.29"N 122°58'47.05"W | Aug, 2013 |  |  | 51 |
| Lynx Hollow Slough | CF17 | Willamette | 176/576 | 491/305 | 43°51'35.07"N 123° 1'24.98"W | Aug, 2013 | 43 | 2013 – 100 bp | 30 |
| Reed Canyon | W17 | Willamette | 29/95 | 190/118 | 45°28'54.66"N 122°37'48.04"W | Jul, 2009 |  |  | 29 |
| Blue Ruin Island | W165 | Willamette | 105/345 | 428/266 | 44°13'35.77"N 123° 9'15.27"W | Nov, 2012 |  |  | 28 |
| Green Island | W173 | Willamette | 111/363 | 441/274 | 44°8'42.02"N 123° 7'4.88"W | Oct, 2012 | 48 | 2013 – 100 bp | 30 |
| Mill Race | W183 | Willamette | 128/420 | 457/284 | 44°2'51.59"N 123° 4'20.22"W | Prior to 2008 |  |  | 29 |
| Science Factory | W184 | Willamette | 125/411 | 459/285 | 44°3'25.50"N 123° 4'30.93"W | 2012, 2013 | 48 | 2013 – 100 bp | 62 |
| Chub Site | W190 | Willamette | 146/478 | 468/291 | 44°1'29.00"N 122°58'29.49"W | Jul, 2012 |  |  | 50 |
| Pudding Creek | W191 | Willamette | 140/460 | 470/292 | 44°1'15.72"N 122°57'37.85"W | Jul, 2012 |  |  | 39 |
| Dougren Slough | W199 | Willamette | 172/565 | 483/300 | 43°58'1.19"N 122°52'8.41"W | Sept, 2013 | 48 | 2013 – 100 bp | 30 |
| Total |  |  |  |  |  |  | 1321 |  | 1419 |

### Collection and scaling of phenotypic data

Measurements and/or counts of 15 morphological traits were collected for all fish in this study including those used in previous work. To capture linear measurements, each fish was photographed with a size standard both laterally and ventrally, and measurements were made using ImageJ (Schneider *et al.,* 2012). Each measurement was performed twice during independent scoring events, and the average of the two scoring events was used for subsequent analysis.

Linear measurements that were taken from lateral photographs are represented in Figure 2A. Standard length (SL) was measured from the rostral-most extent of the upper lip to the posterior end of the caudal peduncle where it meets the caudal fin rays. Body depth was measured from the mid-point of the pelvic joint articulating element to the dorsal outline of the fish. Five linear measurements were taken to capture aspects of defensive traits. Length of the second dorsal and the left pelvic spine were measured from the tip of the spine to the flange at the base of the spine, next to its articulation with basal elements. The height of the ascending process, which extends from the ventral pelvic structure to the lateral plates, was measured from the mid-point of the articulation between the spine and the pelvic plate to the most dorsal tip of the process. In cases where the ascending process ends in multiple cusps, the posterior-most cusp was used. Pelvic structure length was measured from the anterior point of the left anterior process to the caudal tip of the left posterior process (Fig. 2B). Pelvic structure width was taken by measuring the distance between the inner edges of the two pelvic spine joints. Five linear measurements were used to capture variation in head morphology (Fig. 2A). Eye orbit diameter was measured across its widest diameter from a ventral point at the suture between the two suborbital bones. Jaw length was measured from the anterior-most tip of the premaxilla to were the premaxilla and the maxilla meet posteriorly. Dorsal cranial length was measured from the anterior-most tip of the nasal bone to the posterior-most tip of the frontal. The opercle bone roughly resembles a triangle. To capture previously documented differences between adaptive morphologies in this bone we measured from the joint at the dorsoanterior of the opercle to the ventral-most point (JV) and from the joint to the posterior-most point (JP) (Fig. 2A).

**Figure 2.**
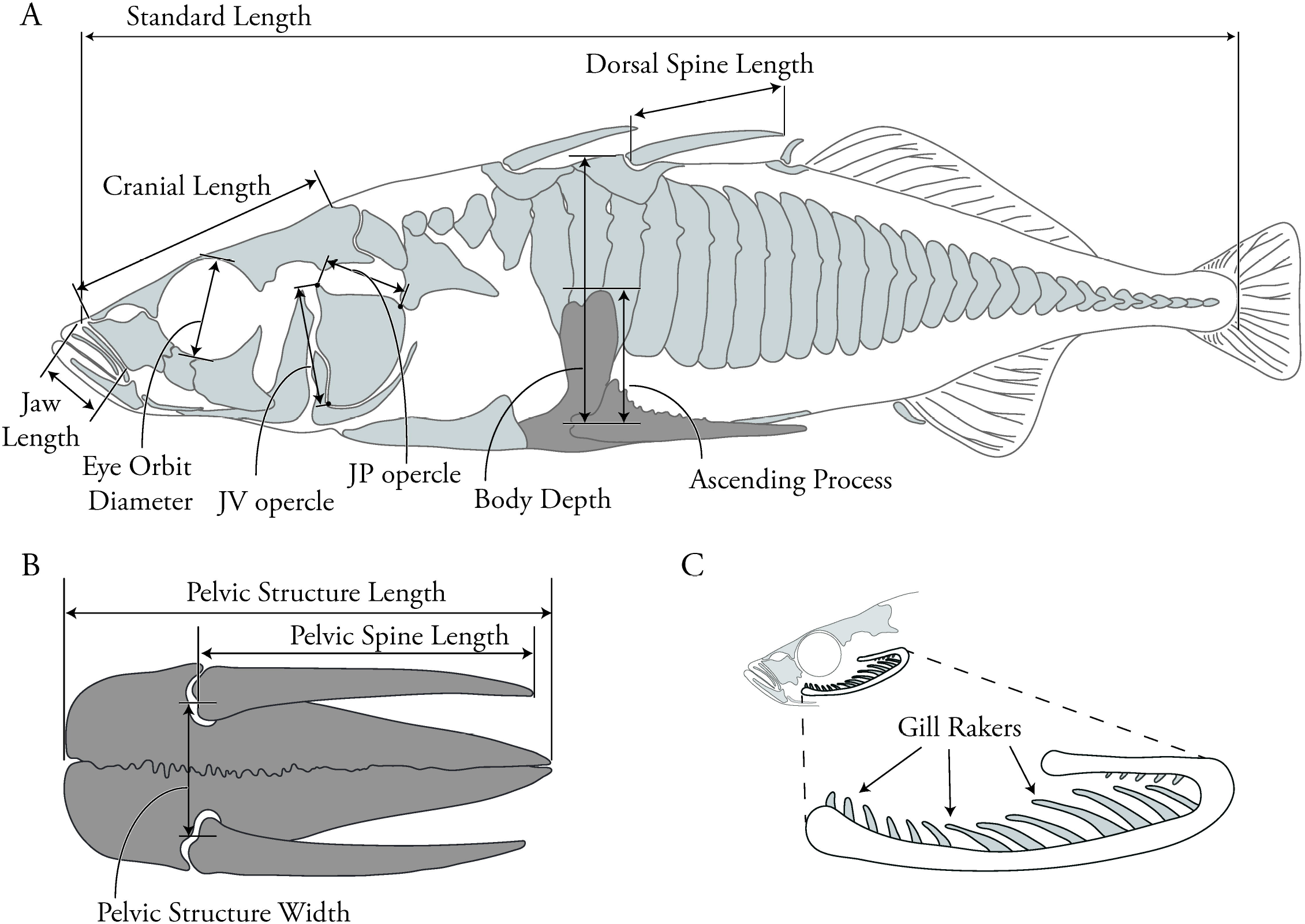
Morphometric measurements selected cover aspects of trophic, defensive, and size traits. (A) Lateral linear measurements: Jaw length, eye orbit diameter, JV opercle, JP opercle, cranial length, body depth, ascending process, pelvic spine length, standard length, and head angle (B) Long gill rakers were counted. (C) Linear measurements of the pelvic structure: pelvic structure length and width, and pelvic spine length. The pelvic structure in A and C has been shaded slightly darker for easier identification.

To size standardize the linear measurements, we employed a method described by Reist (Reist, 1985) that has been used in similar morphological studies (Reimchen & Nosil, 2006; Reimchen *et al.,* 2013). The relationship between measurements of most of the individual traits and SL differed among populations (most, *P* < 0.001 trait x SL interaction, ANCOVA) so population specific slopes for each trait were used to size standardize. In some populations, a significant interaction between some traits and SL was not found: between jaw length and SL in Eel Creek, Middleton Freshwater, and Rabbit Slough, dorsal spine length and SL in Buell-Miller, Finley Swamp, Green Island, Pudding Creek, Meadow Creek, Mud Lake, Bear Paw, and Middleton Fresh, between pelvic spine length and SL in Pudding Creek, Meadow Creek, and Middleton Island, and between eye orbit size, opercle JV/ JP and SL in Rabbit Slough. These traits were mean centered as with the other traits, but not corrected by leaving out the population specific slope term was not included in the equation.

Two meristic traits were also captured: the number of lateral plates on the left side of each fish and the number of long gill rakers from the first arch on the right side (Fig. 2C). These features were counted directly from stained fish on a stereomicroscope. Cranial facial bones were dissected to expose the gill rakers. Because of this destructive sampling the right side rakers were counted to preserve the left side of the head for other phenotypic analyses. Meristic traits were counted twice during independent scoring events and counts that differed were repeated until consistency was reached.

### Analysis of phenotypic variation

To visualize how populations were distributed in phenotypic space, we used Principal Component Analysis (PCA) of the size corrected measurements of individual traits (described above) in R (R Development Core Team, 2013) using the package pcaMethods (Stacklies *et al.,* 2007). Within pcaMethods variables were scaled with unit variance and mean centered before ordination. To test different hypotheses of the partitioning of phenotypic variation between different habitat and regional groupings, Analyses of VAriance (ANOVA) were performed with individual Principle Component (PC) scores using the R package, BaylorEdPsych. Effect size, η^2^, was calculated by dividing the sum of squares treatment by the sum of squares total from the ANOVA analysis.

### Analysis of phenotypic differentiation

To quantify and compare the amount of phenotypic divergence between regional sets of oceanic and freshwater populations we used the measure *P*_ST_ assuming that *c* = *h*^2^ (Storz, 2002; Kaeuffer *et al.,* 2012; Cohen & Dor, 2018). This calculation follows, *P*_ST_ = *c/h*^2^*σ*^2^_*b*_ (*c/h*^2^*σ*^2^_*b*_ + 2*σ*^2^_*w*_). Where *σ*^2^_*b*_ and *σ*^2^_*w*_ are the respective phenotypic variances between and within groups of populations, *c* is an estimate of the proportion of the total variance due to additive genetic effects across populations, and *h*^2^ is heritability, the proportion of phenotypic variance due to additive genetic effects (Brommer, 2011). *P*_ST_ was calculated using the R package Pstat (Da Silva & Da Silva, 2018). Phenotypic PC scores were grouped by habitat type and region (e.g., Alaska oceanic). The *c/h*^2^ default level of 1 was used, 95% confidence intervals were calculated using 1000 bootstrapping data frames. Values of *c/h*^2^ are usually not known for wild populations and can vary across populations so the robustness of this ratio on measures of divergence should be evaluated. To do this we quantified *P*_ST_ with *c/h*^2^ additional values of 0.5 and 0.1.

### RAD library construction and SNP discovery

Restriction site-associated DNA sequencing (RAD-seq) data from previous publications were included: all nine Oregon populations from Catchen et al. (2013) and two populations from Middleton Island, Alaska, one oceanic and one freshwater, from Lescak et al. (2015). New RAD-seq libraries were made for 17 additional populations, using the restriction endonuclease *Sbfl*-HF, and sequenced to 100 or 150 nucleotides on an Illumina HiSeq 2000 or 4000 platform (Table 1), as described in Catchen *et al.* (2013) and Lescak *et al.* (2015). Raw reads were demultiplexed by quality score using the process_radtags program in the *Stacks* software pipeline version 1.46 (Catchen *et al.,* 2011). Processed reads were then aligned against the stickleback genome using GSnap (Wu & Nacu, 2010), allowing for up to five mismatches and gap lengths of two. Only reads with unique alignments were retained and processed through the Stacks pipeline (pstacks, cstacks, sstacks, and populations) to produce a catalog of genotypes and call genotypes for each individual. Only loci that were present in all populations and present in at least 75% of the individuals in each population were used in further analysis.

### Statistical approach

The populations program within the *Stacks* framework was used to calculate genome-wide measures of F_ST_ for each SNP. PCA and STRUCTURE analyses were performed to visualize population grouping and structure of genetic variation. To create a computationally manageable subset of markers from the tens of thousands of RAD loci that were discovered, three sets of ~ 1000 randomly sampled SNPs (restricted to the first SNP/locus) were generated for each collection of individuals included in a comparison (e.g., all Oregon populations or Willamette Basin populations). To investigate and visualize the axes of genetic variation, a PCA was performed on each set of 1000 randomly chosen polymorphic loci using the software Genodive (Meirmans & Van Tienderen, 2004). The mean and standard deviation of PC 1, PC 2, and PC 3 were visualized using R (R Development Core Team, 2013). For all PCAs, a covariance matrix was generated and significance of each PC was tested using a resampling method with 1000 permutations. In all comparisons, all three random SNP sets yielded consistent results (not shown) so results from only one of the sets is presented here. Using the same set of 1000 randomly sampled loci used in the genetic PCAs, STRUCTURE (Pritchard *et al.,* 2000) analyses were performed by running 10 runs for each value of *K* (the number of genetic groupings). Runs for each value of *K* were performed using 40,000 burn-in steps and 40,000 replicates. However, if there was incongruence among the 10 runs using these initial parameters, the number of burn-in steps and the number of replicates for each value of *K* were increased by 10,000 until a congruent answer was reached among all 10 runs (Supplementary Table 1). Results were visualized using Clumpak (Kopelman *et al.,* 2015).

We used Genodive to perform an Analysis of Molecular Variance (AMOVA) (Excoffier *et al.,* 1992) to test the partitioning of genetic variation between different habitat and geographical groupings using the same set of ~ 1000 randomly sampled loci. Coastal and Inland populations were designated by habitat type and not by phenotype. All AMOVAs used an infinite allele model. Significance was determined by resampling with 1000 permutations.

## RESULTS

### Phenotypic variation in Oregon stickleback is partitioned between marine and freshwater habitats and not by geographic proximity

Threespine stickleback were collected throughout northwest Oregon from nearly 40 populations (five oceanic, four coastal freshwater, and 30 inland freshwater) that covered a range of habitat types and geographic regions (Fig. 1, Table 1). Morphometric and meristic data of traits known to diverge between stickleback ecotypes (Fig. 2) were collected from 1283 individuals from these populations and subjected to multivariate analyses. We found that phenotypic variation was predominantly partitioned between populations from marine versus freshwater habitats. The first three axes together explained nearly 70% of the total variation. The major axis of phenotypic variation (PC 1) explained 29% of the total variation and significantly differentiates oceanic and freshwater individuals, (F_1,281_ = 662.8, *P*<2e-16, η^2^ = .34) (Fig. 3A, Supplementary Fig. 1A). The partitioning of phenotypic variation along PC 1 was driven mainly by defensive traits (the number of lateral plates, pelvic girdle width and length, and dorsal and pelvic spine length), and by a trophic trait - the number of gill rakers (Fig. 3B, Table 2). PC 2 explained nearly an equal amount, 25%, of the phenotypic variation and also significantly differentiates oceanic and freshwater individuals, however with a very small effect size, (F_1,281_ = 12.54, *P* < 0.0004, η^2^ = .01) (Fig. 3A, Supplementary Fig. 1A). Variation along PC 2 largely corresponds to craniofacial traits (jaw length, eye orbit diameter, dorsal cranial length, and opercle bone size (Fig. 3B, Table 2). PC 3 explained 13% of the phenotypic variation, significantly partitioned between oceanic and freshwater individuals but with small effect size, (F_1,281_ = 50.5, *P* = 1.92e-12, η^2^ = .04; Supplementary Fig. 1A). Body depth, dorsal cranial length, and pelvic ascending process and girdle length all load highly on PC 3 (Table 2).

**Figure 3.**
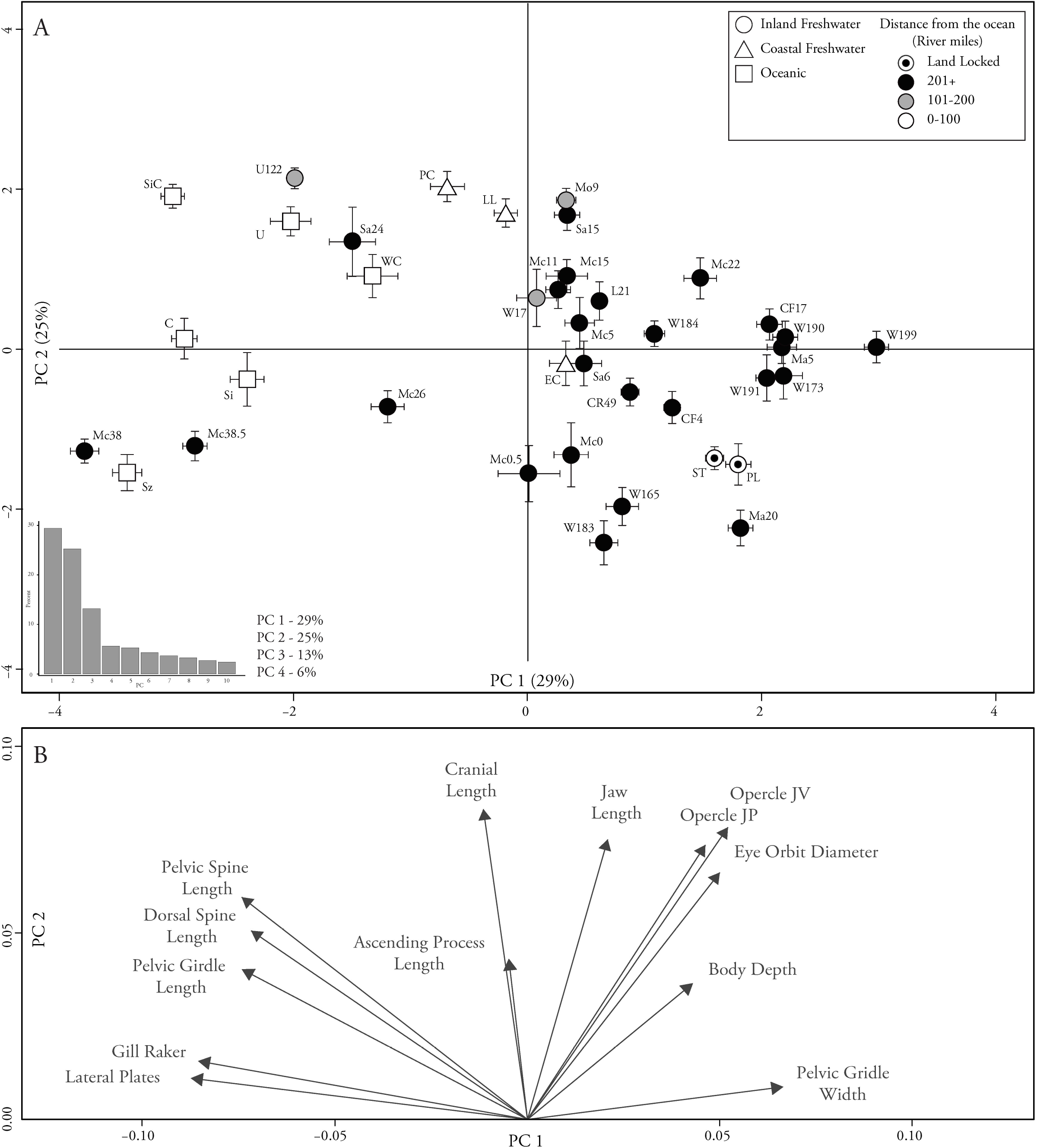
Phenotypic PCA partitions phenotypic variation between oceanic and freshwater stickleback populations. (A) Phenotypic distribution, each symbol represents population level mean PC 1 and 2 scores with bars representing one standard error. PC 1 accounted for 29% of the total phenotypic variation and PC 2 accounted for 25% of the total phenotypic variation. Population symbols have been shaded according to the distance between collection site and the ocean, ranging from light (close) to dark (far) measured in river miles. Circles represent populations found in inland freshwater habitats, triangles represent populations found in coastal freshwater habitats, and squares represent populations found in oceanic habitats. (B) Trait loadings along PC 1 and PC 2. Arrows indicate the direction of a trait’s loading and arrow length indicates its relative strength.

**Table 2.** Trait loadings of the phenotypic PCAs for comparisons of all of the Oregon populations with and without lateral plates and gill rakers and a comparison with Oregon and Alaska populations.

| Trait | PC 1 | PC 2 | PC 3 |
| --- | --- | --- | --- |
| Oregon – All Traits |  |  |  |
| Jaw Length | 0.104 | 0.399 | 0.091 |
| 2<sup>nd</sup> Dorsal Spine | -0.340 | 0.265 | -0.249 |
| Ascending Process | -0.016 | 0.231 | -0.493 |
| Body Depth | 0.231 | 0.170 | -0.359 |
| Left Pelvic Spine | -0.352 | 0.295 | -0.183 |
| Pelvic Girdle Width | 0.322 | 0.061 | -0.433 |
| Pelvic Girdle Length | -0.339 | 0.209 | -0.186 |
| Eye Orbit Diameter | 0.237 | 0.323 | 0.285 |
| Cranial Length | -0.056 | 0.398 | 0.415 |
| Opercle JV | 0.244 | 0.387 | 0.127 |
| Opercle JP | 0.209 | 0.362 | 0.006 |
| Lateral Plates | -0.398 | 0.068 | 0.029 |
| Gill Raker 1<sup>st</sup> Right Arch | -0.397 | 0.074 | 0.181 |
| Oregon - Without Plates and Gill Rakers |  |  |  |
| Jaw Length | 0.414 | 0.004 | 0.082 |
| 2<sup>nd</sup> Dorsal Spine | 0.133 | 0.514 | -0.147 |
| Ascending Process | 0.209 | 0.153 | -0.489 |
| Body Depth | 0.237 | -0.189 | -0.443 |
| Left Pelvic Spine | 0.0159 | 0.532 | -0.070 |
| Pelvic Girdle Width | 0.16 | -0.288 | -0.511 |
| Pelvic Girdle Length | 0.084 | 0.470 | -0.090 |
| Eye Orbit Diameter | 0.389 | -0.200 | 0.241 |
| Cranial Length | 0.364 | 0.135 | 0.441 |
| Opercle JV | 0.448 | -0.149 | 0.097 |
| Opercle JP | 0.410 | -0.096 | -0.011 |
| Oregon vs Alaska - All traits |  |  |  |
| Jaw Length | 0.187 | -0.109 | -0.404 |
| 2<sup>nd</sup> Dorsal Spine | 0.027 | 0.513 | 0.006 |
| Ascending Process | 0.343 | 0.280 | 0.031 |
| Body Depth | 0.393 | 0.103 | 0.172 |
| Left Pelvic Spine | -0.035 | 0.529 | -0.137 |
| Pelvic Girdle Width | 0.420 | 0.091 | 0.245 |
| Pelvic Girdle Length | 0.003 | 0.497 | 0.118 |
| Eye Orbit Diameter | 0.235 | -0.033 | -0.315 |
| Cranial Length | -0.102 | 0.019 | -0.549 |
| Opercle JV | 0.216 | 0.052 | -0.423 |
| Opercle JP | 0.318 | 0.036 | -0.312 |
| Lateral Plates | -0.389 | 0.265 | -0.079 |
| Gill Raker 1<sup>st</sup> Right Arch | -0.388 | 0.157 | -0.174 |

The loss of lateral plates and the reduction of the number of gill rakers are evolutionary phenomena that have been found to be highly associated with the transition from oceanic to freshwater forms in stickleback (Bell & Foster, 1994). Exclusion of these traits had little qualitative effect on the ordination, aside from transposing the first two nearly equal principal components. The first two PCs from this ordination explain similar proportions of the measured variation (30% and 25% respectively) as was found in the analysis with all of the traits. Unlike the analysis that included all of the traits where PC 1 was differentiated between oceanic and freshwater populations, in this reduced analysis it is PC 2 (F_1,281_ 370.7, *P* < 2e-16, η^2^ = .22). The divergence is driven by differences in other defensive traits - dorsal and pelvic spine length, and pelvic girdle length (Supplementary Figs. 1B and 2A, B, Table 2). PC 1 also significantly differentiates oceanic and freshwater individuals, but with little effect size, (F_1,281_ = 6.94, *P* = 0.00852, η^2^ = .005). This PC largely corresponds to differences in craniofacial morphologies - eye orbit diameter, length of dorsal cranial length, and size of the opercle bone (Supplementary Figs. 1B and 2A, B, Table 2). Finally, PC 3 explains 15% of the variation and is driven by differences in the size of the ascending process, body depth, pelvic width and dorsal cranial length. This component also significantly differentiates oceanic and freshwater individuals but with little effect size (F_1,281_ = 87.19, *P* < 2e-16, η^2^ = .06) (Supplementary Fig. 1B).

### Partitioning of genetic variation is qualitatively different from partitioning of phenotypic variation in Oregon stickleback

RAD sequencing data was generated for 1321 individuals from 15 Oregon and 2 Alaska populations ranging from 12 to 48 fish per population (Table 1), and these new data sets were augmented with previously published data from 9 Oregon (Catchen *et al.,* 2013) and two Middleton Island, Alaska populations (Lescak *et al.,* 2015). After stringent quality filtering and processing with the Stacks pipeline ~ 20,000 SNPs were identified.

Unlike phenotypic variation, which we found to be partitioned chiefly by marine versus freshwater habitat types, genotypic variation in Oregon is partitioned across geographic regions. Analysis of genetic variation including all of the Oregon populations differentiates coastal versus inland populations, both in multivariate ordination using PCA (Fig. 4) and in STRUCTURE analysis (Fig. 5). In our multivariate ordination, PC 1 accounts for the majority (~ 63%) of the total genetic variation and clearly separates coastal from inland populations (Fig. 4, Table 3). In addition, some separation occurs between coastal freshwater and coastal oceanic populations along PC 2, though this outcome is not statistically significant. STRUCTURE analysis using the lowest level of clustering, *K* = 2, produces qualitatively similar genetic partitioning as seen in the genetic PCA. Clusters are anchored by the same two geographic regions, with both marine and freshwater coastal populations clustering to the exclusion of inland populations (Fig. 5). An interesting set of exceptions for both the genetic, and phenotypic, analyses are the populations from 1) Milk Creek (Mo9) (a tributary of the Molalla River), 2) Buell-Miller Slough (Sa24) of the Santiam River, 3) Riverbend (Mc11), Walterville (Mc26), and Leaburg (Mc38) (all three from the McKenzie River), and 4) Page Road from the Umpqua River (U122) (Fig 3 & 4).

**Figure 4.**
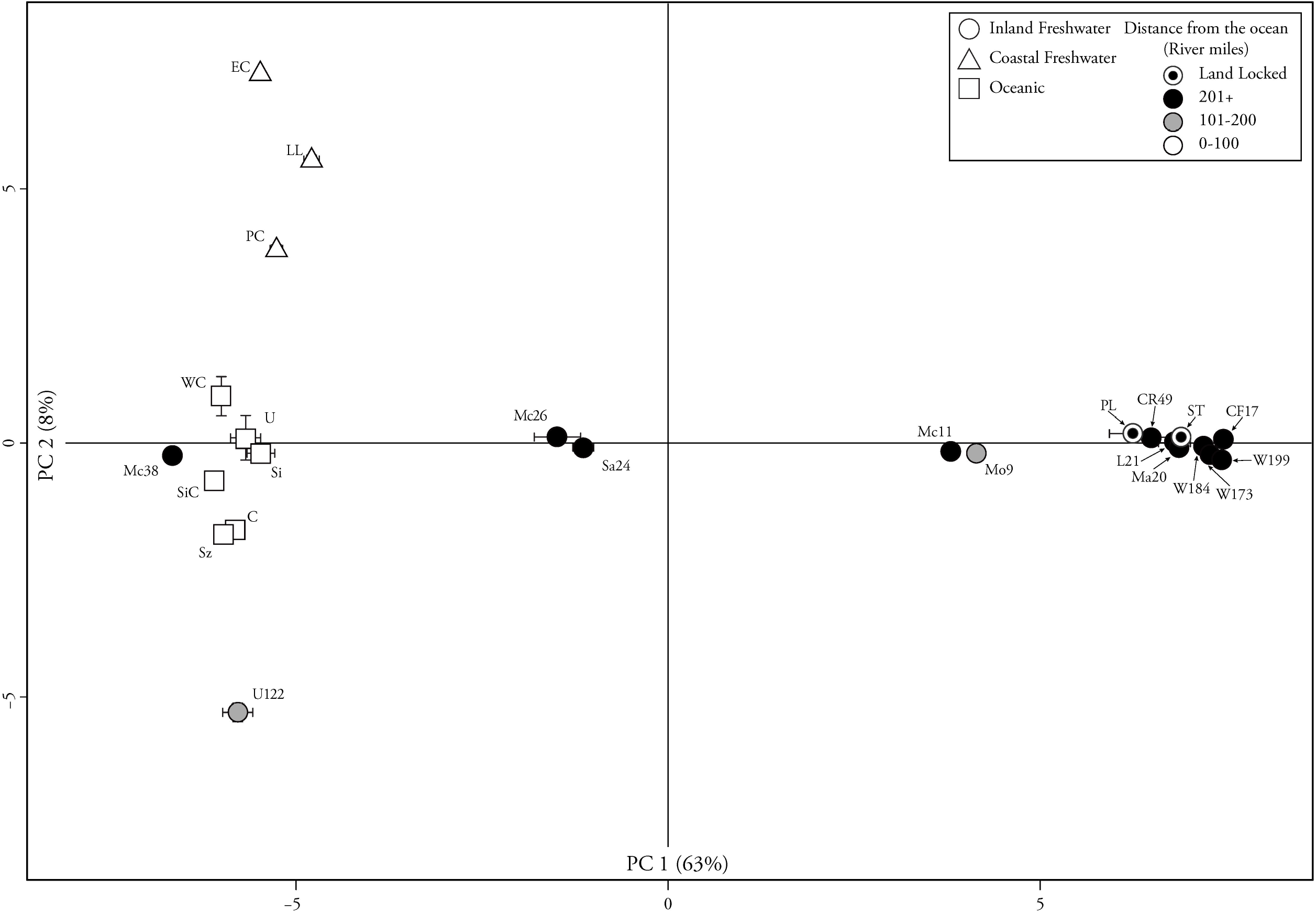
Genetic variation in Oregon stickleback is partitioned between regions and not by habitat types. PC 1 explains 63% of the genetic variation and separates coastal populations on the left and inland populations on the right. The average PC 1 and PC 2 score of each population is plotted, with bars for one standard error. Population symbols have been shaded according to the distance between collection site and the ocean, ranging from light (close) to dark (far) measured in river miles. Circles represent populations found in inland freshwater habitats, triangles are populations found in coastal freshwater habitats, and squares are populations found in oceanic habitats.

**Figure 5.**
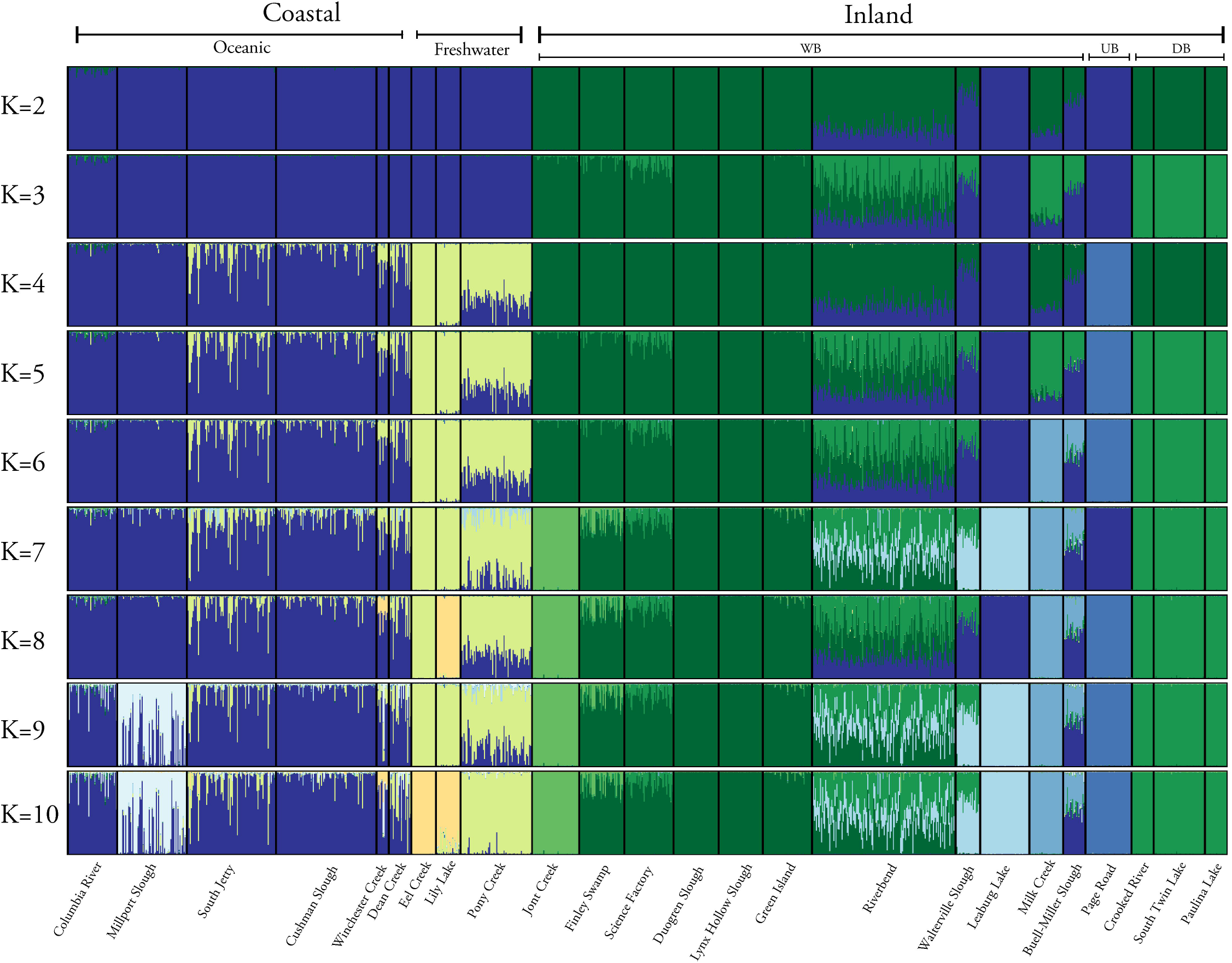
Oregon populations of stickleback are genetically structured by region, but with exceptions. Each vertical bar represents an individual with colors representing the posterior probability of membership to that group, as estimated by STRUCTURE. Each level of *K* represents 10 repeats that were run with increasing levels of burn-in and repetitions until congruence was reached. In general, at low levels of *K*, populations are partitioned between coastal and inland regions. With increasing levels of *K* the two regions increasingly are partitioned by location. However, some inland populations (e.g., Leaburg Mc38) do not strictly follow these patterns.

**Table 3.** Genetic PCA results that included 20 Oregon populations. Principle components 1-6 with eigenvalues, percent variance explained, cumulative variance explained, and p-value of each PC. The first PC accounts for the vast majority of the genetic variation and is the only axis that is statistically significantly.

| PC | Eigenvalue | % Variance | Cumulative | *P*-value |
| --- | --- | --- | --- | --- |
| 1 | 12.810 | 63.03 | 63.03 | 0.01 |
| 2 | 1.647 | 8.10 | 71.13 | 0.7 |
| 3 | 1.285 | 6.323 | 80.81 | 0.53 |
| 4 | 0.681 | 3.35 | 84.05 | 1 |
| 5 | 0.660 | 3.25 | 86.44 | 1 |
| 6 | 0.484 | 2.38 | 88.39 | 1 |

At increasing levels of *K* the two geographic regions that correspond to coastal and inland remain intact, however these two clusters become increasingly subdivided into groupings that correspond to habitat type and geographic location. At *K* = 3, the inland cluster is partitioned between the Willamette and Deschutes basins, which are separated by the Cascade mountain range (Fig 5). The Deschutes Basin populations show affinity with some Willamette Basin populations (e.g., Riverbend and Science Factory) consistent with the hypothesis that the upper Deschutes Basin populations stem from a recent introduction of fish originating in the Willamette Basin (Yake, 2003; Davis *et al.,* 2010; Catchen *et al.,* 2013). At *K* = 4, the coastal cluster splits by habitat into oceanic and freshwater groupings. However partial membership in both coastal clusters by some individuals in each habitat could be an indication of recent gene flow or incomplete partitioning of genetic variation. One interesting observation is the splitting off and subsequent re-clustering of some populations with increasing values of K, e.g., the behavior of Central Oregon populations when *K* = 3-5. We interpret this as most likely due to the fine partitioning of genetic variation in some of the populations, warranting more sensitive analytical approaches. At *K* = 5-10 regional and habitat groups are increasingly subdivided. Coastal populations retain membership in respective oceanic and freshwater groupings, but the coastal freshwater populations become subdivided (e.g., Lily Lake splits off at *K* = 8). In the inland cluster the populations Riverbend, Walterville, Leaburg, Milk Creek, Buell-Miller, and Page Road, that at *K* = 2-3 demonstrate affinity with the coastal cluster, then increasingly separate from the coast and one another at higher *K* values. Finally, at the largest levels of K, inland populations begin to disband, with Jont creek breaking into a unique group at *K* = 7.

This pattern of the partitioning of genetic variation is also present when comparing average measures of population differentiation and analysis of molecular variance (AMOVA). On average, FST values between oceanic and inland populations (excluding the inland populations with outlying genetic affinity to oceanic populations) was ~ 0.16, with a range of 0.12 (Cushman Slough vs Finley Grey Creek Swamp) to 0.25 (Dean Creek vs Dougren Slough) (Supplementary Table 2). This is similar to previously reported levels of divergence between oceanic and Willamette Basin populations (Catchen *et al.,* 2013). In contrast stickleback populations found in Oregon’s coastal freshwater habitats demonstrated the lowest average genome wide divergence from oceanic populations (average coastal freshwater compared to average oceanic, F_ST_ = 0.06). This measure of divergence is similar to what we found between young oceanic and freshwater pairs from southcentral Alaska (F_ST_ = 0.056) and Middleton Island (F_ST_ = 0.057). Finally, we used AMOVA to quantify how genetic variation is partitioned within and between individuals, populations, and geographical regions (coastal and inland). We found that most of the variation was partitioned between individuals (~ 46%) however there was little partitioning of variation between individuals within the populations (~ 4%). Similar to the patterns we found using PCA and STRUCTURE, we found significant variation partitioned between the populations within each geographical region (~ 23%) and significant variation partitioned between the coastal and inland geographical regions (~ 28%) (Table 4).

**Table 4.** AMOVA table demonstrating the amount of partitioning of genetic variation within and between individuals, among populations nested within coastal and inland groupings, and among coastal and inland groupings. This analysis shows that the majority of variation not accounted for by individuals or populations is partitioned between coastal and inland populations.

| Source of Variation | Nested in | % Variance | $F$-stat | $P$-value |
| --- | --- | --- | --- | --- |
| Within Individual | ---- | 0.459 | $F_{IT}$ | ---- |
| Among Individual | Population | 0.040 | $F_{IS}$ | 0.001 |
| Among Population | Coastal vs Inland | 0.225 | $F_{SC}$ | 0.001 |
| Among Coastal vs Inland | ---- | 0.276 | $F_{CT}$ | 0.001 |

### Phenotypic variation in Oregon stickleback is partitioned in a similar manner as in Alaska populations but is accompanied by much greater genetic divergence

How phenotypic and genetic variation is partitioned in young postglacial stickleback populations has been intensively pursued (e.g., Schluter, 1993; Bell & Orti, 1994; Cresko *et al.,* 2004; Shapiro *et al.,* 2004; Colosimo *et al.,* 2005; Kimmel *et al.,* 2005; Hohenlohe *et al.,* 2010; Jones *et al.,* 2012b; Catchen *et al.,* 2013; Reimchen *et al.,* 2013; Bassham *et al.,* 2018). An open question is how well these patterns extend to stickleback populations that have inhabited much older landscapes. To address this question, we compared patterns of the partitioning of phenotypic and genetic variation that we found in Oregon to younger stickleback systems such as in southcentral Alaska (~ 12,000 to 15,000 years old) and Middleton Island (~ 55 years old).

Using our findings from the STRUCTURE and genetic PC analyses we selected Oregon populations that exhibit a strong partitioning of genetic variation between oceanic and freshwater habitats to use in a phenotypic analysis comparing these diverging Oregon populations with oceanic and freshwater divergent populations located in south central and Middleton Island Alaska. Using PCA we found similar patterns of phenotypic divergence in Oregon and Alaska populations along the major axis and quantified and compared this divergence with measures of *P*_ST_. PC 1, which explained 30% of the variation, groups oceanic populations, both Alaska and Oregon (Fig. 6A). This PC significantly differentiates oceanic and freshwater populations regardless of geographic region, with a very large effect size, (F_662_ = 1273.8, *P* < 2e-16, η^2^ = 0.58; Fig. 6, Supplementary Fig. 3). Both systems demonstrated nearly the same amounts of phenotypic divergence along PC 1 with *P*_ST_ values of 0.997 and 0.995 between oceanic and freshwater groupings in the Oregon and Alaska populations respectively (Supplementary Table 3). Because we did not know the values of *c/h*^2^ for these populations, different values were used to determine the robustness of the measures of divergence that we found. These measures appear to be robust over reduced values of *c/h*^2^, values as low as 0.1 produced very similar results, *P*_ST_ of PC 1= 0.975 and 0.954 in the Oregon and Alaska systems (Supplementary Table 3). The phenotypic divergence between the oceanic and freshwater groupings was driven by the number of lateral plates and gill rakers, depth of the ascending process and body, eye orbit diameter, and size of the opercle (Fig 6B, Table 2). Interestingly, dorsal and pelvic spine length and pelvic girdle length did not weigh as heavily on this axis of differentiation as it did in the analysis that included only the Oregon populations. Somewhat unexpectedly, we did find significant regional partitioning of phenotypic variation of PC 1 between Oregon and Alaska regardless of habitat type, with a moderate effect size (F_662_ = 233.17, *P* < 2e-16, η^2^ = 0.26).

**Figure 6.**
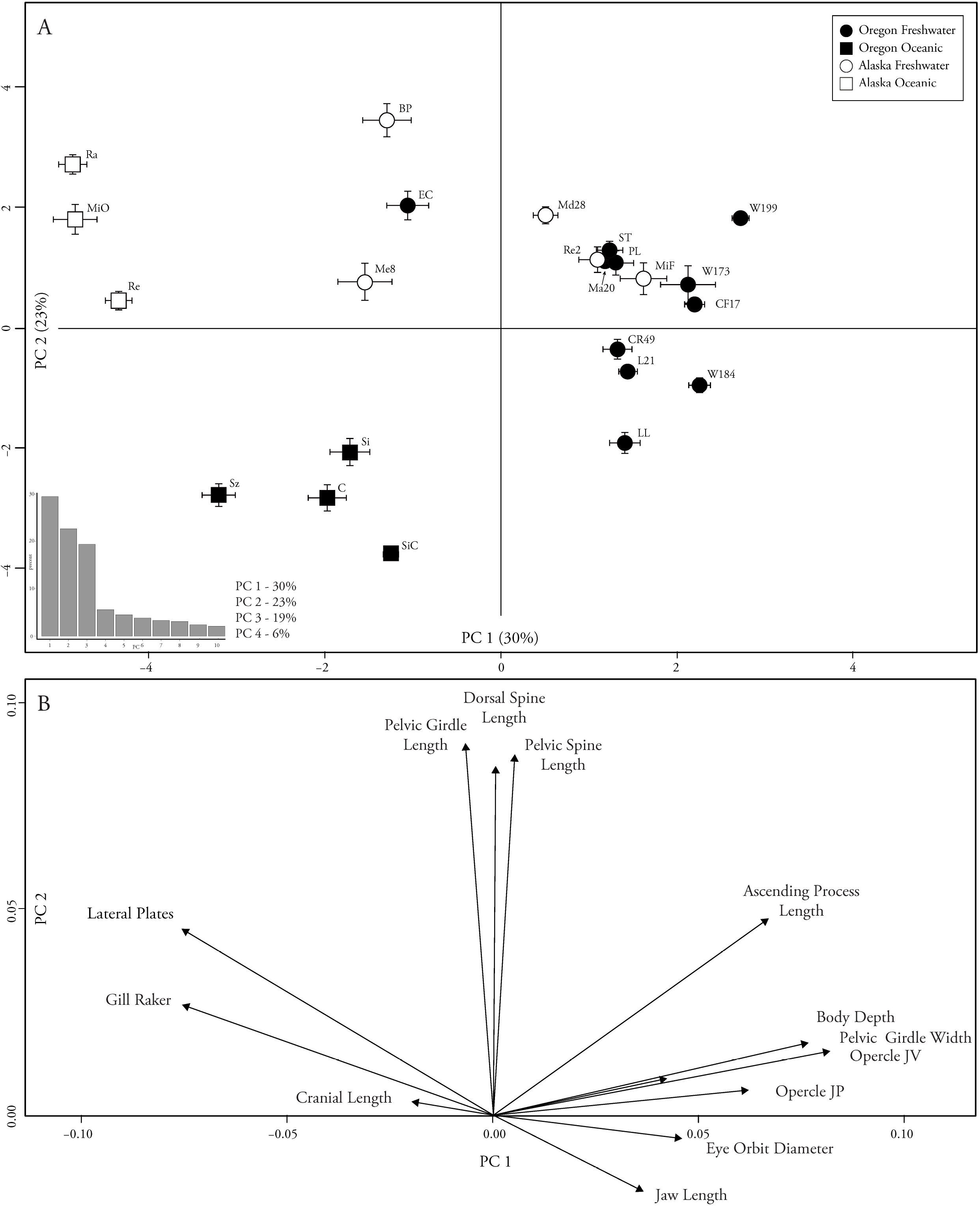
Oregon and Alaska stickleback populations show similar phenotypic divergence between ecotypes. Phenotypic variation is partitioned between oceanic and freshwater ecotypes along PC 1 which explains 30% of the variation, oceanic populations to the left and freshwater populations to the right, squares versus circles. PC 2 explains 23% of the variation. Population means are plotted, with bars for one standard error of the populations mean. Black symbols represent Oregon populations and white symbols are Alaska. Circles represent populations found in freshwater habitats and squares are populations found in oceanic habitats.

PC 2 also partitioned phenotypic variation between habitat and regional groupings. PC 2, which explained 23% of the total variation, significantly partitioned variation by habitat, (F_662_ = 300.7, *P* <2e-16, η^2^ = 0.31) and region, (F_662_ = 334.9, *P* <2e-16, η^2^ = 0.33) with both groupings having similar effect sizes (Fig. 6. Supplementary Fig. 3). As in PC 1, phenotypic variation is clearly partitioned between Oregon oceanic and freshwater populations in PC 2, with these groups demonstrating a large measure of phenotypic divergence, *P*_ST_ = 0.996. However, unlike PC 1, there is less partitioning of phenotypic variation between Alaska habitat types, with a very small measure of phenotypic divergence, *P*_ST_ = 0.082 (Fig. 6, Supplementary Table 3). PC 3 also significantly differentiates phenotypic variation between habitat and by region but with small effect size, (F_662_ = 10.65, *P* = 0.00116, η^2^ = .02) and (F_662_ = 28.21, *P* = 1.49e-07, η^2^ = .04) (Supplementary Fig 3).

It is interesting to note that there is clear separation of Oregon and Alaska oceanic forms along PC 1 and 2 (Fig 6). To investigate further, an additional PC analysis was performed with only the oceanic populations from these two regions. In this analysis phenotypic variation is significantly partitioned between Oregon and Alaska along PC 1 (F_208_ = 598.4, *P* <2e-16, η^2^ = 0.74) (Supplementary Fig. 4A). This is driven by differences in dorsal and pelvic spine length, pelvic width and length, and the length of the ascending process and body depth (Supplementary Fig. 4B). Phenotypic differences in oceanic populations from different parts of their range have been documented before (Defaveri & Merila, 2013; Morris *et al.,* 2018) however this is counter to the idea that marine stickleback consist largely of a single phenotypic form (Bell & Foster, 1994), and implies a greater diversity of marine populations throughout this species’ global range is likely.

Two phenotypically divergent pairs, one from Middleton Island and another from southcentral Alaska and the divergent populations in Oregon were compared to assess genomic levels of differentiation between these systems. PCA was again used. We found that PC 1, which accounts for ~ 51% of the partitioning of genetic variation and which is the only significant PC (Table 5), again separates Oregon populations between coastal and inland geographic regions regardless of habitat type, just as in the PCA including only Oregon populations (Fig. 7). Surprisingly, all of the Alaska populations, from both systems and ecotypes, cluster with Oregon coastal populations (Fig. 7). Likewise, genome-wide average F_ST_ between Alaska oceanic and freshwater populations is ~ 0.07, similar to the divergence between Oregon oceanic and Oregon coastal freshwater populations (F_ST_ ~ 0.06). By comparison, Oregon inland freshwater populations are more than twice as diverged from Oregon oceanic, with an average F_ST_ of ~ 0.16.

**Figure 7.**
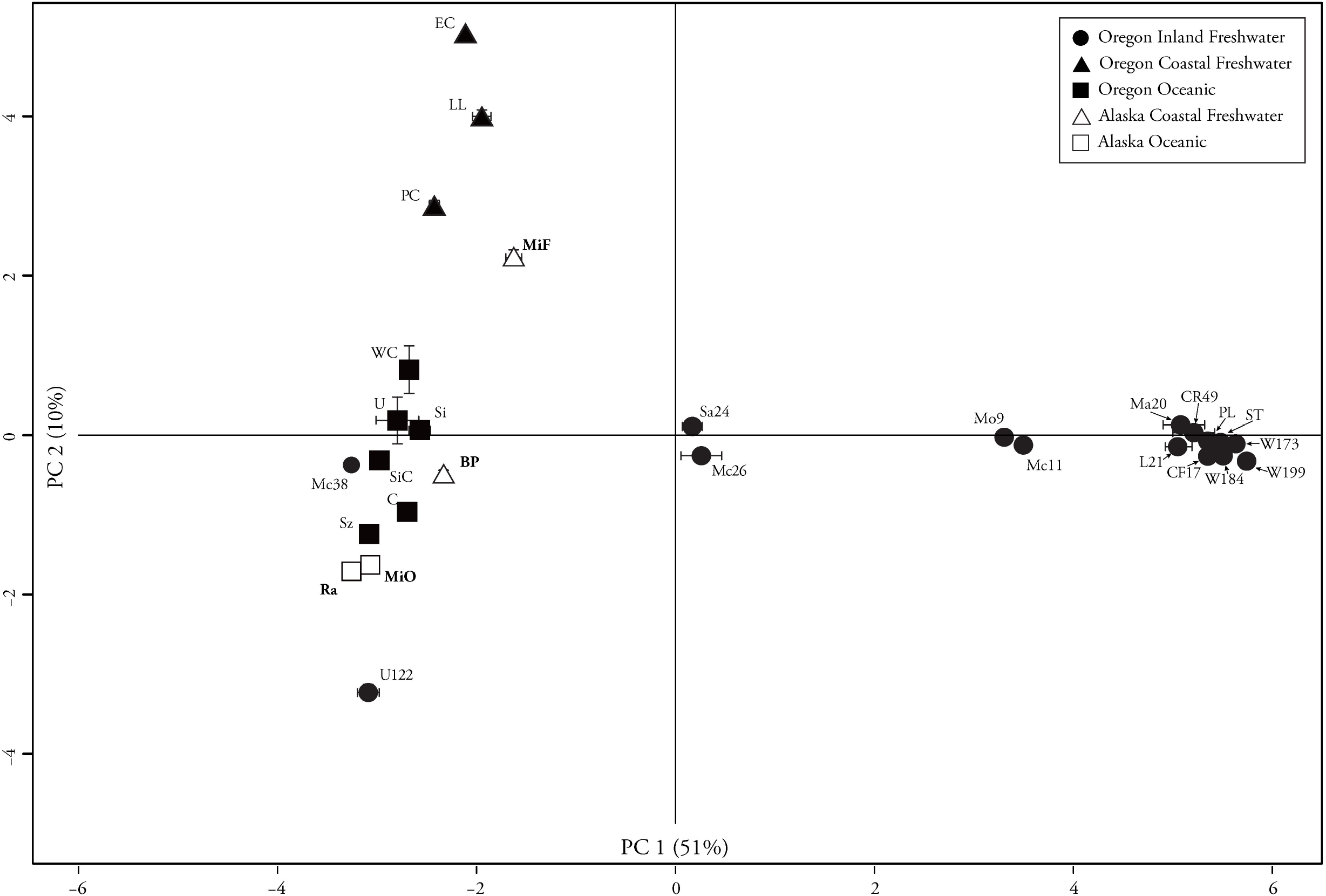
Alaska populations group with Oregon coastal populations in genetic PCA. The only significant principal component (PC1) explained 51% of the genetic variation. The average PC 1 and PC 2 scores of each population are plotted, with bars for one standard error of the mean. Circles represent populations found in inland freshwater habitats, triangles are populations found in coastal freshwater habitats, and squares are populations found in oceanic habitats. Black symbols represent Oregon populations and white symbols are Alaska. Alaska populations labels are in bold.

**Table 5.** Genetic PCA results that included Oregon and Alaska populations. Principle components 1-6 with eigenvalues, percent variance explained, cumulative variance explained, and p-value of each PC. The first PC accounts for the vast majority of the genetic variation and is the only axis that is statistically significantly.

| PC | Eigenvalue | % Variance | Cumulative | *P*-value |
| --- | --- | --- | --- | --- |
| 1 | 13.23 | 50.86 | 50.86 | 0.02 |
| 2 | 2.66 | 10.22 | 61.08 | 0.38 |
| 3 | 1.64 | 6.3 | 67.38 | 0.86 |
| 4 | 1.4 | 5.4 | 72.78 | 0.80 |
| 5 | 1.07 | 4.12 | 76.9 | 0.91 |
| 6 | 0.89 | 3.41 | 80.31 | 0.97 |

### Varying degrees of genetic divergence exist between populations within and between Oregon’s watersheds

Dougren Slough, Lynx Hollow, and Jont Creek are all located far upstream within their individual drainages, 12, 17.5, and 21 river miles respectively. Science Factory and Green Island are located downstream from Dougren Slough and Lynx Hollow along the valley floor, in the main stem of the Willamette near the confluence of these drainages (Fig. 1). Finley Grey Creek Swamp is also located on the valley floor in a location that likely has high connectivity with the main stem via historical and present-day flooding (Benner & Sedell, 1997). There is strong genetic divergence among the upstream populations, with F_ST_ values ranging from 0.33 (Jont creek versus Lynx Hollow) to 0.37 (Dougren Slough versus Lynx Hollow) and less divergence between these populations and their downstream counterparts, with F_ST_ values ranging from 0.09 (Dougren Slough versus Science Factory) to 0.19 (Lynx Hollow versus Finley Swamp) (Supplementary Table 2).

While genome-wide divergence was strikingly high between low and high elevation drainage populations, we observed extraordinarily high within-species genetic divergence among stickleback populations from the Willamette, Umpqua, and Deschutes watersheds. These values of F_ST_ ranged from 0.51 (Page Road (Umpqua) and Science Factory (Willamette), and Page Road and South Twin (Deschutes)) to 0.61 (Page Road and Dougren Slough (Willamette)) (Supplementary Table 2). Such extreme divergence has been reported between two *Gasterosteus* species, *G. aculeatus and G. nipponicus,* estimated to have diverged 0.68-1 million years ago (Ravinet *et al.,* 2018).

### Inland freshwater populations are genetically structured by drainage, with exceptions

To better understand the patterns of genetic divergence within the Willamette Basin, we focused multivariate genetic analyses on populations collected throughout this large basin. In an analysis including eleven populations from tributaries of the Willamette River, and from the Willamette itself, the first and the only significant genetic PC accounted for ~ 64% of the variation. Populations along the PC 1 axis were differentiated into groups with low versus higher than average lateral plate counts (Fig. 8, Table 6). In STRUCTURE at *K* = 2, populations are again clustered into low and extra-plated groups (Fig 9A). However, Leaburg stickleback, from a relatively high elevation in the McKenzie river, form a discrete cluster. Jont Creek, Finley Swamp, Science Factory, Dougren Slough, Lynx Hollow, and Green Island, all located on the valley floor, form the other cluster. Interestingly, populations geographically located between Leaburg and the valley floor group are a mixture of both clusters. At *K* = 3-5, the northern Willamette Basin extra-plated Milk Creek and Buell-Miller populations form an exclusive cluster, with the Buell-Miller population having partial membership coefficients in the Leaburg cluster (Fig. 9A). These findings were confirmed using AMOVA, whereby a significant amount of variation was partitioned between extra- and low-plated populations (~ 16 %) that was not partitioned within individuals (~ 54%) or between populations (~ 28%) (Table 7). Further analysis of only the low plated populations resulted in these populations being more partitioned at higher values of *K.* Dougren Slough, Lynx Hollow Slough, and Jont Creek all separate into nearly three discrete freshwater clusters at *K* = 3, and Finley Grey Creek Swamp, Green Island, and Science Factory all harbor mixtures of genotypes from these three discrete clusters (Fig. 9B).

**Figure 8.**
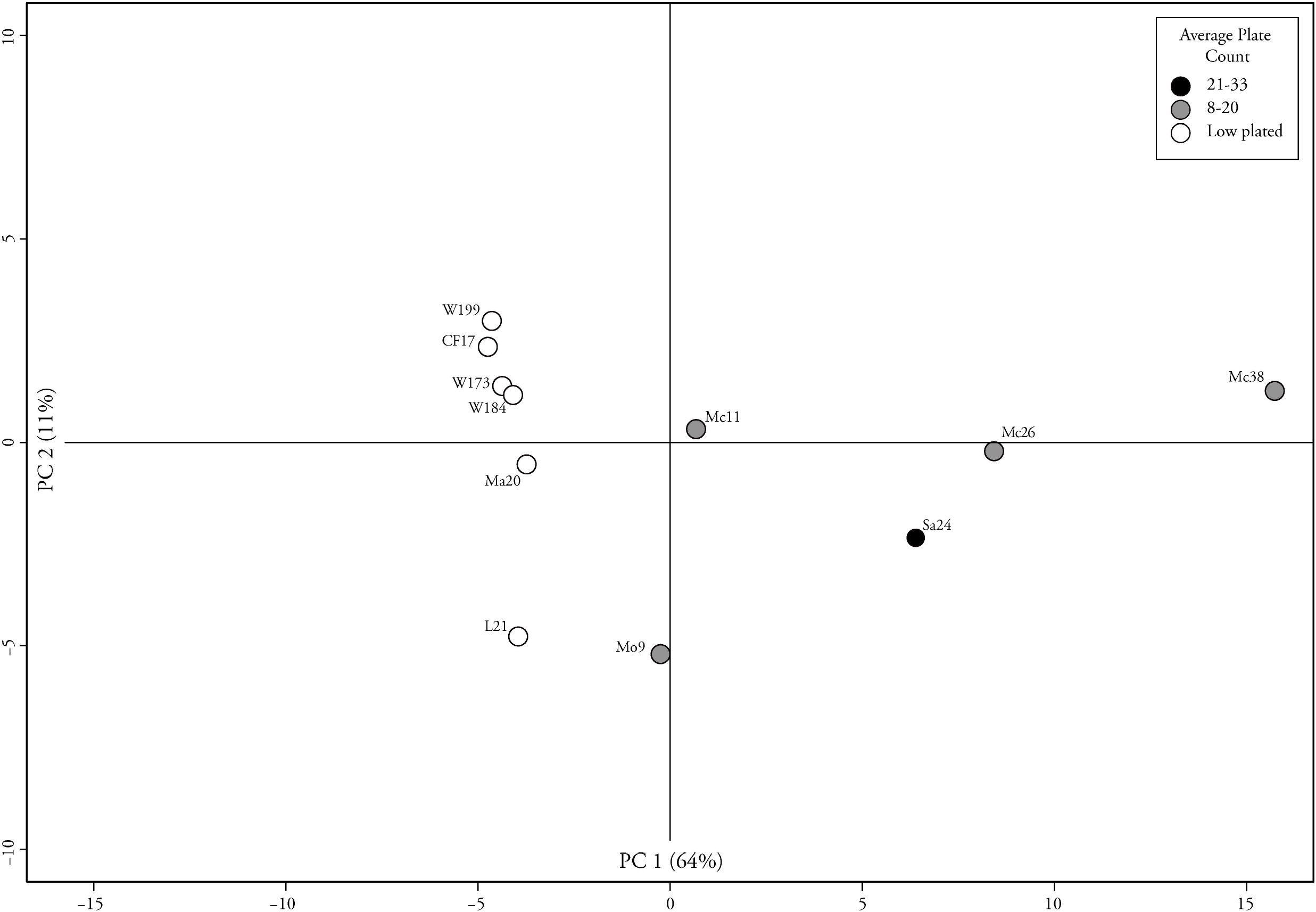
Genetic variation is partitioned between low and extra-plated populations. The only significant principal component (PC1) explained 64% of the genetic variation and partitions variation between fish with few lateral plates and those with average lateral plate counts between 8 and 33. The average PC 1 and PC 2 scores of each population are plotted, with bars for one standard error. Symbols are color coded for the average number of plates of each population.

**Figure 9.**
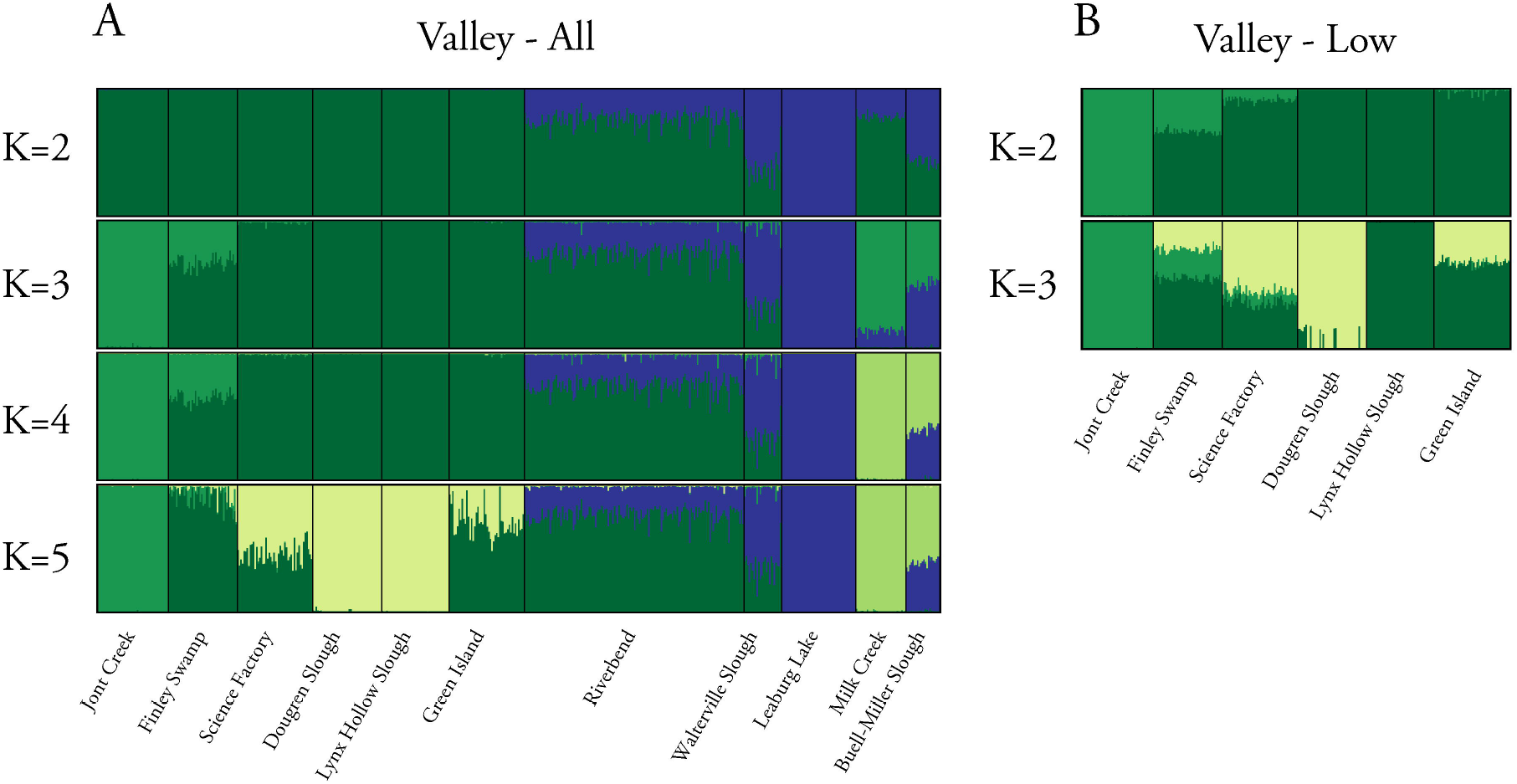
Willamette Basin populations are genetically structured by drainage, despite connectivity of habitats. STRUCTURE groupings are plotted for multiple levels of *K.* Each vertical bar represents an individual color-coded by its posterior probability of membership to a group. (A) Extra and low plated populations. At low levels of *K* populations are partitioned between average low and extra plated populations with increased partitioning with increasing levels of *K.* (B) Low plated populations. At *K* = 3, Jont Creek, Dougren Slough, and Lynx Hollow all group independently while populations downstream display individuals with mixed memberships.

**Table 6.** Genetic PCA results that included Willamette Basin populations. Principle components 1-6 with eigenvalues, percent variance explained, cumulative variance explained, and p-value of each PC. The first PC accounts for the vast majority of the genetic variation and is the only axis that is statistically significantly.

| PC | Eigenvalue | % Variance | Cumulative | *P*-value |
| --- | --- | --- | --- | --- |
| 1 | 48.33 | 63.8 | 56.37 | 0.03 |
| 2 | 8.012 | 10.56 | 74.36 | 0.88 |
| 3 | 5.037 | 6.64 | 81 | 0.96 |
| 4 | 3.738 | 4.93 | 85.9 | 0.99 |
| 5 | 3.514 | 4.63 | 90.6 | 0.99 |
| 6 | 2.619 | 3.45 | 94.0 | 0.98 |

**Table 7.** AMOVA table demonstrating the amount of partitioning of genetic variation within and between individuals, among populations nested within low and extra-plated groupings, and among low and extra-plated groupings.

| Source of Variation | Nested in | % Variance | *F*-stat | *P*-value |
| --- | --- | --- | --- | --- |
| Within Individual | ----- | 0.535 | *F*<sub>IT</sub> | ---- |
| Among Individual | Population | 0.027 | *F*<sub>IS</sub> | 0.001 |
| Among Population | Low vs Extra | 0.277 | *F*<sub>SC</sub> | 0.001 |
| Among Low vs Extra | ----- | 0.162 | *F*<sub>CT</sub> | 0.003 |

The structuring of the low plated populations and the pattern of divergence that we identified is consistent with observations in other riverine fish systems, which show strong unidirectional gene flow in upstream populations presumably resulting from increasing river gradients with gain in elevation (Hernandezmartich & Smith, 1990; Shaw *et al.,* 1991; Castric *et al.,* 2001), (Fig. 10). However, the extra- plated populations did not fit this pattern. The discovery of populations that appear to have genetic and phenotypic affinity to oceanic populations that were collected very far from the coast was unexpected and warranted further investigation.

**Figure 10.**
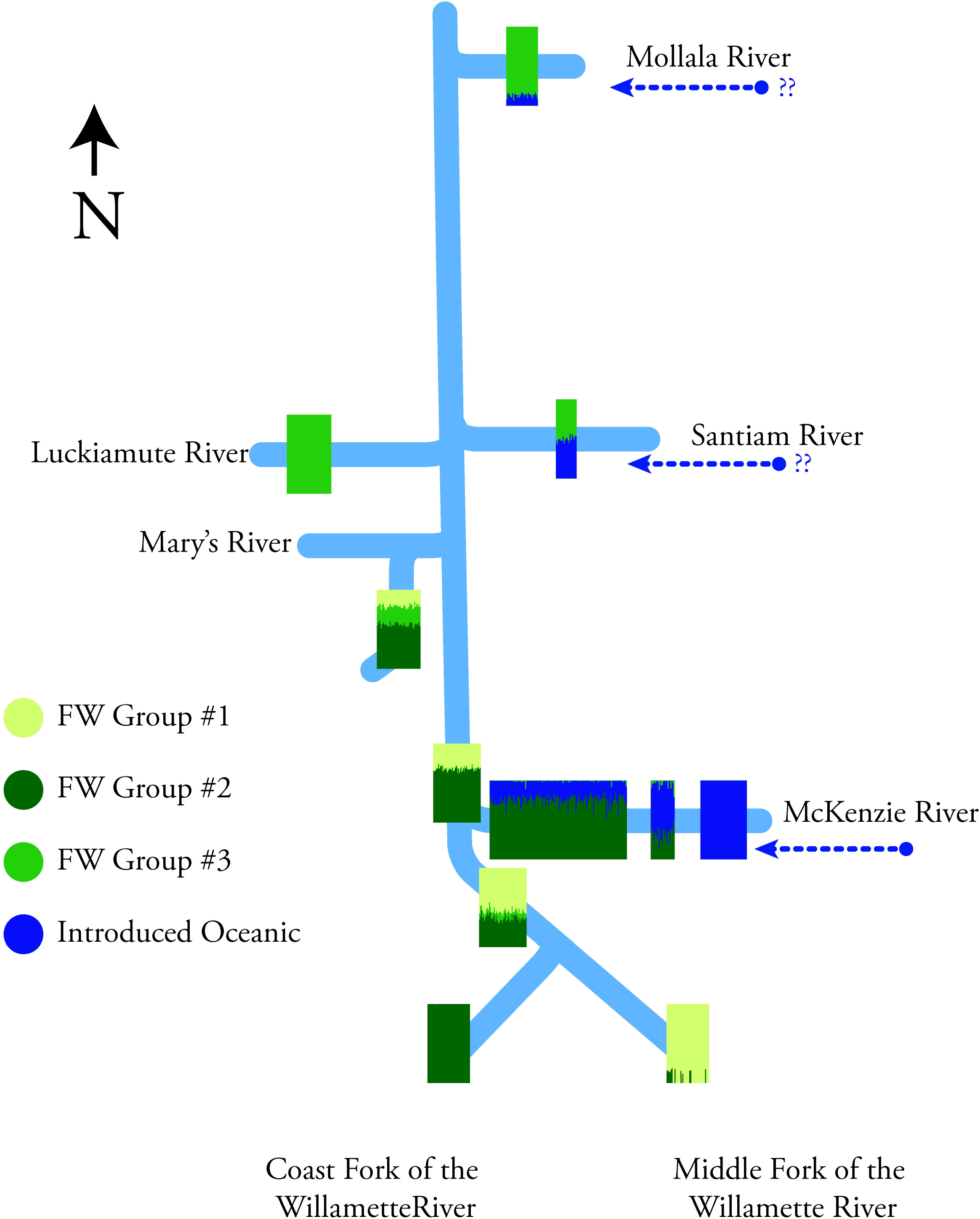
Genetic structure of Willamette Basin populations emulates the pattern of distance, directional flow, and sequence of confluences, but with exceptions. The genetic clusters found with STRUCTURE are plotted on a representation of the major rivers of the basin. Distinct clusters (group 1 and 2) in the upper forks of the Willamette’s main stem appear to be almost fully partitioned between Coast and Middle Forks. Downstream, these constituents grade into a third distinct genetic group (group 3), which appears to predominate lower in the watershed. Oceanic genotypes (blue) are differentially introgressed into freshwater genotypes along three rivers flowing down the west slope of the Cascade foothills.

### Some Oregon inland populations cluster phenotypically and genetically with oceanic populations

One startling finding was the presence of extra and high plated stickleback in rivers deep in the Willamette and Umpqua Basins. Phenotypic analysis including multiple traits places these populations squarely within oceanic groupings or in phenotypically intermediate groups (Fig. 3A). Excluding lateral plates and gill rakers from analyses, these populations still cluster with the oceanic populations or with phenotypically intermediate groups along PC 2, which in this analysis separated oceanic and freshwater forms (Supplementary Fig. 2A).

Genetic analysis supports and clarifies these relationships. Along PC 1, Leaburg Lake (Mc38) (McKenzie River) and Page Road (U122) (Umpqua River) cluster distinctly with the coastal populations (and within the oceanic populations along PC 2) (Fig. 4). Populations from Riverbend (Mc11) and Walterville (Mc26) (both of the McKenzie River), Buell-Miller Slough (Sa24) (Santiam River), and Milk Creek (Mo9) (Molalla River) are genetic intermediates between coastal populations and inland low plated populations. At all levels of *K* tested in STRUCTURE analysis there is partial membership of these ocean-like freshwater populations with marine populations. This is best exemplified at *K* = 2-8, where Leaburg, and to a lesser extent, Page Road, Milk Creek, and Buell-Miller, partially or fully cluster with marine populations (Fig. 5). Consistent with STRUCTURE patterns, ocean-like populations in the Willamette Basin display less genome-wide divergence from marine fish (average F_ST_ ~ 0.08) than they do from other Willamette Basin populations (average F_ST_ ~ 0.21) (Supplementary Table 2). This modest degree of divergence between the oceanic-like Willamette Basin and actual oceanic populations is similar to genetic divergence reported between oceanic and very young freshwater populations on Middleton Island (average F_ST_ ~ 0.067; (Lescak *et al.,* 2015).

## DISCUSSION

### Stickleback inhabiting old freshwater landscapes harbor a similar degree of morphological divergence from marine fish but greater genetic divergence than those inhabiting younger freshwater habitats

Freshwater stickleback across western Oregon display many aspects of stereotypical phenotypic divergence from their marine relatives that has been noted throughout the range of this species (Bell & Foster, 1994; Conte *et al.,* 2015). This “freshwater syndrome” includes reduction in the number of lateral plates and gill rakers, the reduction in the size of aspects of the pelvic structure and pelvic spines, and the reduction in the size of the dorsal spines. We did not find, however, populations with characteristics that have been encountered more rarely, as partial or complete loss of the pelvic structure that has been documented in Alaska and California (Bell, 1987), nor did we find evidence for a major partitioning of phenotypic variation within freshwater habitats that has been occasionally documented, for example between the benthic and limnetic stickleback species pairs in British Columbia (Mcphail, 1984). The well-studied loss of lateral plates and reduction of gill rakers are major factors we observed in the partitioning of phenotypic space, as variation in both of these traits loads heavily along the vector distinguishing oceanic and freshwater ecotypes. When excluding lateral plates and gill rakers from the analysis, we find that other measured morphological variation - encompassing a suite of defensive and trophic traits - are partitioned between Oregon oceanic and freshwater populations to a similar extent as it is in comparisons of oceanic with freshwater populations that were founded thousands or just dozens of years ago (Bell & Foster, 1994; Cresko *et al.,* 2004; Lescak *et al.,* 2015).

Similarities of divergence in Oregon with stickleback populations in post-glacial landscapes fade when we explore population genomic metrics. Surprisingly, the major partitioning of genetic variation in Oregon stickleback is between coastal (both marine and freshwater) and inland populations, which is out of alignment with phenotypic and life history affinities. Importantly, this pattern of divergence is in stark contrast to other stickleback systems in which the major axis of phenotypic and genetic divergence is between oceanic and freshwater populations. The discordance between the phenotypic and genetic data in the Oregon populations argues for several important findings. First, the small amount of genetic divergence between coastal populations - regardless of habitat - suggests that coastal freshwater habitats were founded relatively recently, and that these freshwater populations could experience ongoing gene flow with oceanic fish. Second, the degree of genetic differentiation of inland versus coastal populations is consistent with the hypothesis that Willamette Basin populations predate the last glacial maximum and/or are isolated by distance.

The observation of much more similar patterns of phenotypic divergence than of genetic divergence among very young (50 years on Middleton Island), postglacial (southcentral Alaska), and much older (inland Oregon) freshwater populations argues that the “freshwater syndrome” evolves rapidly and hinges on reuse of standing genetic variation across particular genomic loci (Barrett & Schluter, 2008; Hohenlohe *et al.,* 2010; Feulner *et al.,* 2013; Terekhanova *et al.,* 2014; Marques *et al.,* 2016; Bassham *et al.,* 2018). However, this rapid burst of phenotypic and corresponding genetic change may then be followed by long periods of phenotypic evolutionary stasis underlain by continuing neutral genetic divergence. If some of the Oregon freshwater populations do indeed predate the last glacial maximum, as evidence presented here suggests, a testable hypothesis is that these populations might harbor much older freshwater haplotypes at genomic loci implicated in re-use during recent transitions to freshwater (Bassham *et al.,* 2018; Nelson & Cresko, 2018).

### An unexpected finding of oceanic fish far inland in major Oregon watersheds

Despite the overall pattern of oceanic and freshwater phenotypic divergence, and coastal and inland population genomic divergence, an anomalous phenotypic and genetic clustering of some inland Oregon populations with coastal groups was at first confusing. These inland oceanic-like populations were not localized to a specific geographic region, but were found in the higher elevation extent of the species’ range in two major watersheds and in four major rivers within these watersheds. Others have reported extra-plated populations in and around Oregon (Rutter, 1896; Hagen & Gilbertson, 1972), northern California (Baumgartner & Bell, 1984), and in inland regions of Europe and British Columbia (Münzing, 1963; Reimchen *et al.,* 2013). The presence of these morphologies far inland presents different possible scenarios. These patterns could be the result of recent or episodic introductions, perhaps by humans, of coastal fish into these systems. Alternatively, selection could have favored the maintenance or reiteration of oceanic phenotypes, as has been reported in stickleback from British Columbia (Reimchen *et al.,* 2013) and Washington State (Kitano *et al.,* 2008).

Suggestive of human introduction of marine fish, stickleback were not found in Leaburg Lake in the 1980’s despite intensive sampling at Leaburg Dam by wildlife regulators (Zakel, 1984). Analysis of genetic divergence shows that the current Leaburg population is more closely related to oceanic populations than it is to the low-plated fish found throughout the Willamette Basin. Strikingly, Leaburg is an order of magnitude more diverged from a population just 39 miles away in same river (F_ST_ = 0.38) than it is from oceanic fish ~ 480 river miles away (F_ST_ = 0.039).

Favoring a hypothesis of long-term maintenance of oceanic morphological traits in at least one inland locale, on the other hand, is a historical survey from 1896. This study reported that the fourteen stickleback collected approximately 120 river miles from the sea in the north Umpqua were fully plated (Rutter, 1896). We found that stickleback near this location today are likewise fully plated (average plate count of 32) and are phenotypically like marine fish in other traits as well. Despite these morphological affinities, this inland population is more than twice as genetically diverged from marine fish (F_ST_ = 0.11), than are other inland oceanic-like populations, such as Buell-Miller (F_ST_ = 0.049) on the Santiam River and Leaburg Lake on the McKenzie (F_ST_ = 0.058). Presumably, this Umpqua River population has maintained a phenotypically oceanic shape for at least 120 years even while it has been genomically becoming distinct.

Here we present a comprehensive phenotypic and genetic analysis of dozens of populations of threespine stickleback from diverse habitats and geographic regions from across Oregon. This rich dataset reveals a complex panorama of degrees of natural divergence and signatures of anthropogenic introductions. Young freshwater populations in coastal lakes and streams showed strong genetic affinity to oceanic populations that inhabit the estuaries and bays along the coast, despite their phenotypic divergence from those marine fish and their phenotypic congruence with older, isolated freshwater populations located far inland. Some aspects of the mosaic described here echo what has been reported in other stickleback systems, but surprising discoveries in the Oregon system add new colors and textures, and extend the impact that studies of this small fish have made to our understanding of how ecological and evolutionary processes play out across landscapes.

## ACKNOWLEDGEMENTS

We thank Steven James, Shira Mali, Savanah Olroyd, Erik Parker, Aimee Schultz, Roberta Torunsky, Jack Peplinski, and Mikaeli Dirling for help with collection of phenotypic data. We also thank Brian Bangs, Paul Olmstead, Sara Akins, Brian Cannon, Randy Wildman, and Laurie Weitkamp from Oregon Department of Fish and Wildlife (ODFW) and National Oceanic and Atmospheric Administration (NOAA) for help with procurement of many of the stickleback samples used in this manuscript. This research was supported primarily by National Science Foundation 0949053 and IOS 102728 (both to W.A.C.). Additional support came from National Institutes of Health grant 1R24GM079486-01A1 and OD011199 (both to W.A.C.).

## CONFLICTS OF INTEREST

The authors declare no conflicts of interest.

---

**Supplementary Figure 1.** Significant phenotypic differences between ecotypes in Oregon threespine stickleback shown by box plots of PCs 1-3. (A) All phenotypic traits included. (B). All traits except the number of lateral plates and gill rakers.

**Supplementary Figure 2.** Phenotypic PCA of Oregon populations excluding lateral plate and gill raker counts. (A) Phenotypic PCA partitions phenotypic variation between oceanic and freshwater stickleback populations along PC 2. (B) Dorsal and pelvic spine length and pelvic girdle length and width drive these differences.

**Supplementary Figure 3.** Box plots demonstrating significant difference between oceanic and freshwater ecotypes and between regions among Oregon and Alaska threespine stickleback populations in PC 1-3.

**Supplementary Figure 4.** PCA of phenotypic traits comparing Oregon and Alaska oceanic populations.

