## Supplemental Table 1 for "Genetic divergence outpaces phenotypic divergence among threespine stickleback populations in old freshwater habitats"

|  | Oregon | | Willamette Basin  All | | Willamette Basin  Low | |
| --- | --- | --- | --- | --- | --- | --- |
| *K* | Burn-in  /Reps | LnProb | Burn-in  /Reps | LnProb | Burn-in  /Reps | LnProb |
| 2 | 40K/40K | -203064.6 | 80K/80K | -192327.4 | 40K/40K | -146127.1 |
| 3 | 40K/40K | -199004.8 | 80K/80K | -182609.8 | 40K/40K | -128470.4 |
| 4 | 100K/100K | -182458.8 | 120K/120K | -174225.8 | 40K/40K | -114729.8 |
| 5 | 100K/100K | -178469.0 | 120K/120K | -168926.5 | 40K/40K | -107323.9 |
| 6 | 40K/40K | -176297.7 | 120K/120K | -166026.9 | - | - |
| 7 | 40K/40K | -176677.1 | - | - | - | - |
| 8 | 60K/60K | -171951.1 | - | - | - | - |
| 9 | 60K/60K | -166699.6 | - | - | - | - |
| 10 | 60K/60K | -163546.4 | - | - | - | - |

**Supplementary Table 1.** STRUCTURE run statistics for Oregon, Willamette Basin, and Willamette Basin low runs. For each value of the number of burn-in steps and repetitions are reported along with the LnProb value.
