## Supplemental Table 2 for "Genetic divergence outpaces phenotypic divergence among threespine stickleback populations in old freshwater habitats"

|  | Sz | Si | SiC | U | WC | EC | LL | PC | U122 | Mo9 | Sa24 | L21 | Ma20 | W184 | W199 | CF17 | W173 | Mc11 | Mc26 | Mc38 | CR | ST | PL | Ra1 | BP | MiF | MiO |
| --- | --- | --- | --- | --- | --- | --- | --- | --- | --- | --- | --- | --- | --- | --- | --- | --- | --- | --- | --- | --- | --- | --- | --- | --- | --- | --- | --- |
| CR | 0.016 | 0.012 | 0.011 | 0.026 | 0.022 | 0.090 | 0.056 | 0.045 | 0.098 | 0.109 | 0.044 | 0.152 | 0.142 | 0.142 | 0.158 | 0.155 | 0.147 | 0.081 | 0.052 | 0.057 | 0.128 | 0.143 | 0.128 | 0.016 | 0.071 | 0.066 | 0.016 |
| Sz |  | 0.014 | 0.012 | 0.029 | 0.024 | 0.097 | 0.059 | 0.051 | 0.102 | 0.123 | 0.053 | 0.168 | 0.157 | 0.157 | 0.174 | 0.172 | 0.163 | 0.095 | 0.061 | 0.058 | 0.143 | 0.158 | 0.143 | 0.020 | 0.077 | 0.075 | 0.021 |
| Si |  |  | 0.005 | 0.016 | 0.009 | 0.058 | 0.036 | 0.028 | 0.077 | 0.090 | 0.037 | 0.126 | 0.118 | 0.118 | 0.128 | 0.127 | 0.123 | 0.079 | 0.041 | 0.040 | 0.104 | 0.120 | 0.104 | 0.018 | 0.059 | 0.052 | 0.017 |
| SiC |  |  |  | 0.017 | 0.011 | 0.062 | 0.039 | 0.033 | 0.074 | 0.091 | 0.037 | 0.126 | 0.118 | 0.118 | 0.128 | 0.127 | 0.123 | 0.081 | 0.042 | 0.039 | 0.104 | 0.120 | 0.104 | 0.017 | 0.058 | 0.054 | 0.016 |
| U |  |  |  |  | 0.030 | 0.115 | 0.057 | 0.043 | 0.164 | 0.159 | 0.062 | 0.234 | 0.212 | 0.210 | 0.246 | 0.242 | 0.218 | 0.082 | 0.076 | 0.084 | 0.210 | 0.212 | 0.217 | 0.031 | 0.100 | 0.076 | 0.037 |
| WC |  |  |  |  |  | 0.097 | 0.049 | 0.031 | 0.148 | 0.148 | 0.058 | 0.213 | 0.193 | 0.192 | 0.221 | 0.218 | 0.201 | 0.085 | 0.067 | 0.073 | 0.185 | 0.195 | 0.188 | 0.029 | 0.091 | 0.070 | 0.032 |
| EC |  |  |  |  |  |  | 0.069 | 0.085 | 0.330 | 0.299 | 0.151 | 0.406 | 0.366 | 0.364 | 0.431 | 0.424 | 0.375 | 0.171 | 0.175 | 0.202 | 0.362 | 0.367 | 0.377 | 0.090 | 0.204 | 0.143 | 0.105 |
| LL |  |  |  |  |  |  |  | 0.045 | 0.161 | 0.146 | 0.080 | 0.193 | 0.178 | 0.177 | 0.200 | 0.197 | 0.184 | 0.096 | 0.087 | 0.096 | 0.170 | 0.178 | 0.172 | 0.059 | 0.107 | 0.076 | 0.065 |
| PC |  |  |  |  |  |  |  |  | 0.162 | 0.165 | 0.077 | 0.222 | 0.206 | 0.206 | 0.228 | 0.225 | 0.213 | 0.131 | 0.086 | 0.095 | 0.186 | 0.208 | 0.187 | 0.048 | 0.111 | 0.086 | 0.051 |
| U122 |  |  |  |  |  |  |  |  |  | 0.424 | 0.212 | 0.580 | 0.509 | 0.505 | 0.609 | 0.604 | 0.515 | 0.230 | 0.265 | 0.281 | 0.528 | 0.508 | 0.561 | 0.085 | 0.252 | 0.218 | 0.106 |
| Mo9 |  |  |  |  |  |  |  |  |  |  | 0.107 | 0.194 | 0.160 | 0.159 | 0.237 | 0.230 | 0.174 | 0.061 | 0.158 | 0.301 | 0.146 | 0.149 | 0.164 | 0.105 | 0.244 | 0.195 | 0.124 |
| Sa24 |  |  |  |  |  |  |  |  |  |  |  | 0.186 | 0.162 | 0.161 | 0.201 | 0.199 | 0.172 | 0.056 | 0.059 | 0.128 | 0.148 | 0.161 | 0.154 | 0.050 | 0.124 | 0.098 | 0.057 |
| L21 |  |  |  |  |  |  |  |  |  |  |  |  | 0.163 | 0.175 | 0.341 | 0.329 | 0.171 | 0.070 | 0.248 | 0.416 | 0.218 | 0.194 | 0.267 | 0.142 | 0.325 | 0.265 | 0.171 |
| Ma20 |  |  |  |  |  |  |  |  |  |  |  |  |  | 0.088 | 0.180 | 0.192 | 0.086 | 0.048 | 0.215 | 0.370 | 0.143 | 0.138 | 0.165 | 0.132 | 0.293 | 0.242 | 0.158 |
| W184 |  |  |  |  |  |  |  |  |  |  |  |  |  |  | 0.095 | 0.135 | 0.089 | 0.045 | 0.214 | 0.367 | 0.118 | 0.114 | 0.141 | 0.132 | 0.292 | 0.240 | 0.157 |
| W199 |  |  |  |  |  |  |  |  |  |  |  |  |  |  |  | 0.371 | 0.160 | 0.065 | 0.268 | 0.430 | 0.299 | 0.248 | 0.348 | 0.145 | 0.336 | 0.273 | 0.176 |
| CF17 |  |  |  |  |  |  |  |  |  |  |  |  |  |  |  |  | 0.157 | 0.067 | 0.263 | 0.429 | 0.294 | 0.251 | 0.356 | 0.144 | 0.333 | 0.272 | 0.174 |
| W173 |  |  |  |  |  |  |  |  |  |  |  |  |  |  |  |  |  | 0.045 | 0.230 | 0.382 | 0.161 | 0.152 | 0.186 | 0.137 | 0.301 | 0.249 | 0.163 |
| Mc11 |  |  |  |  |  |  |  |  |  |  |  |  |  |  |  |  |  |  | 0.047 | 0.172 | 0.032 | 0.048 | 0.035 | 0.079 | 0.156 | 0.141 | 0.082 |
| Mc26 |  |  |  |  |  |  |  |  |  |  |  |  |  |  |  |  |  |  |  | 0.106 | 0.186 | 0.210 | 0.193 | 0.058 | 0.153 | 0.117 | 0.066 |
| Mc38 |  |  |  |  |  |  |  |  |  |  |  |  |  |  |  |  |  |  |  |  | 0.358 | 0.376 | 0.373 | 0.056 | 0.178 | 0.151 | 0.065 |
| CR |  |  |  |  |  |  |  |  |  |  |  |  |  |  |  |  |  |  |  |  |  | 0.023 | 0.085 | 0.123 | 0.284 | 0.222 | 0.150 |
| ST |  |  |  |  |  |  |  |  |  |  |  |  |  |  |  |  |  |  |  |  |  |  | 0.058 | 0.133 | 0.296 | 0.244 | 0.159 |
| PL |  |  |  |  |  |  |  |  |  |  |  |  |  |  |  |  |  |  |  |  |  |  |  | 0.123 | 0.291 | 0.226 | 0.152 |
| Ra1 |  |  |  |  |  |  |  |  |  |  |  |  |  |  |  |  |  |  |  |  |  |  |  |  | 0.063 | 0.062 | 0.008 |
| BP |  |  |  |  |  |  |  |  |  |  |  |  |  |  |  |  |  |  |  |  |  |  |  |  |  | 0.125 | 0.073 |
| MiF |  |  |  |  |  |  |  |  |  |  |  |  |  |  |  |  |  |  |  |  |  |  |  |  |  |  | 0.068 |

**Supplementary Table 2.** Genome-wide averaged F_ST_ for all pairwise comparisons of populations included in the genotypic analysis. Abbreviations for populations can be found in table 1.
