## Supplemental Table 3 for "Genetic divergence outpaces phenotypic divergence among threespine stickleback populations in old freshwater habitats"

|  |  | P_ST_ Values | 95% Low CI | 95% Up CI |
| --- | --- | --- | --- | --- |
| c/h = 1.0 |  |  |  |  |
| Alaska | PCA_axis_1 | 0.99516238 | 0.994425308 | 0.995878412 |
|  | PCA_axis_2 | 0.08211634 | 0.012397382 | 0.685417655 |
|  | PCA_axis_3 | 0.91779211 | 0.837206302 | 0.965406804 |
|  | PCA_axis_4 | 0.65895524 | 0.170203214 | 0.897379817 |
| Oregon | PCA_axis_1 | 0.99742158 | 0.996779655 | 0.997817546 |
|  | PCA_axis_2 | 0.9957181 | 0.994328026 | 0.996584435 |
|  | PCA_axis_3 | 0.02759286 | 0.018543678 | 0.685318023 |
|  | PCA_axis_4 | 0.90907914 | 0.865008157 | 0.932521624 |
| c/h = 0.5 |  |  |  |  |
| Alaska | PCA_axis_1 | 0.99037134 | 0.986770363 | 0.992931815 |
|  | PCA_axis_2 | 0.04281612 | 0.000258995 | 0.576483997 |
|  | PCA_axis_3 | 0.84807376 | 0.61111054 | 0.924382582 |
|  | PCA_axis_4 | 0.49137453 | 0.014208166 | 0.794690948 |
| Oregon | PCA_axis_1 | 0.99485642 | 0.993472581 | 0.99597674 |
|  | PCA_axis_2 | 0.99147272 | 0.988945735 | 0.993234423 |
|  | PCA_axis_3 | 0.01398943 | 0.000253391 | 0.570634774 |
|  | PCA_axis_4 | 0.83331356 | 0.568924347 | 0.920063358 |
| c/h = 0.1 |  |  |  |  |
| Alaska | PCA_axis_1 | 0.95364216 | 0.937391128 | 0.966510976 |
|  | PCA_axis_2 | 0.00886694 | 5.98E-05 | 0.197614801 |
|  | PCA_axis_3 | 0.52750584 | 0.243415779 | 0.705398836 |
|  | PCA_axis_4 | 0.16192923 | 0.001590739 | 0.447914925 |
| Oregon | PCA_axis_1 | 0.97480056 | 0.968698741 | 0.979852673 |
|  | PCA_axis_2 | 0.95876992 | 0.948440745 | 0.96724023 |
|  | PCA_axis_3 | 0.00282955 | 3.30E-05 | 0.216731089 |
|  | PCA_axis_4 | 0.49996441 | 0.195404155 | 0.70906937 |

**Supplementary Table 3.** *P*_ST_ measures, with 95% confidence intervals, using the first four phenotypic PC scores comparing oceanic and freshwater regional sets. Three different values for c/h were used to test robustness of the ratio.
