## Supplementary figures and images for "Genetic divergence outpaces phenotypic divergence among threespine stickleback populations in old freshwater habitats"

### Supplemental Fig. 1

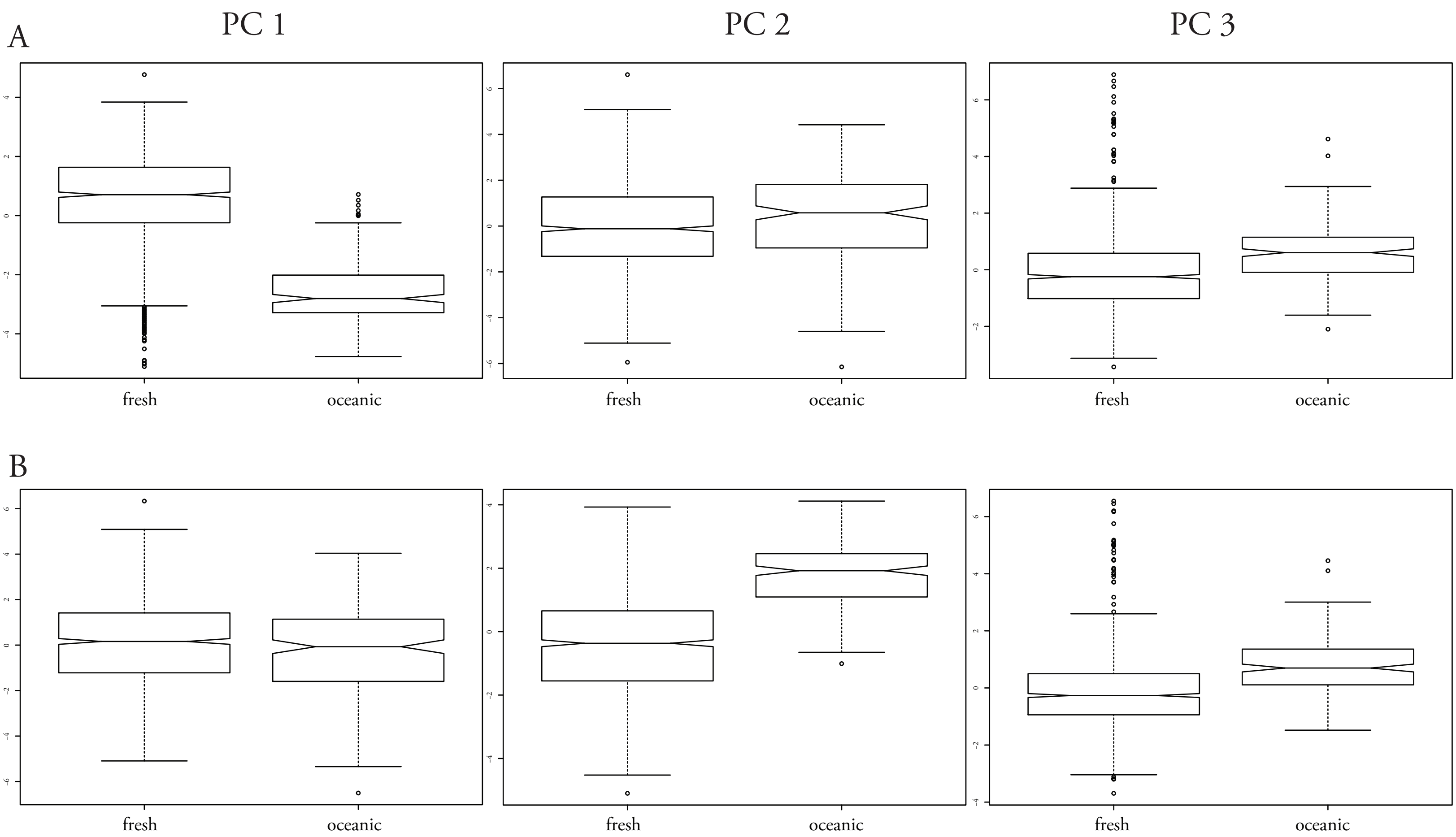

### Supplemental Fig. 2

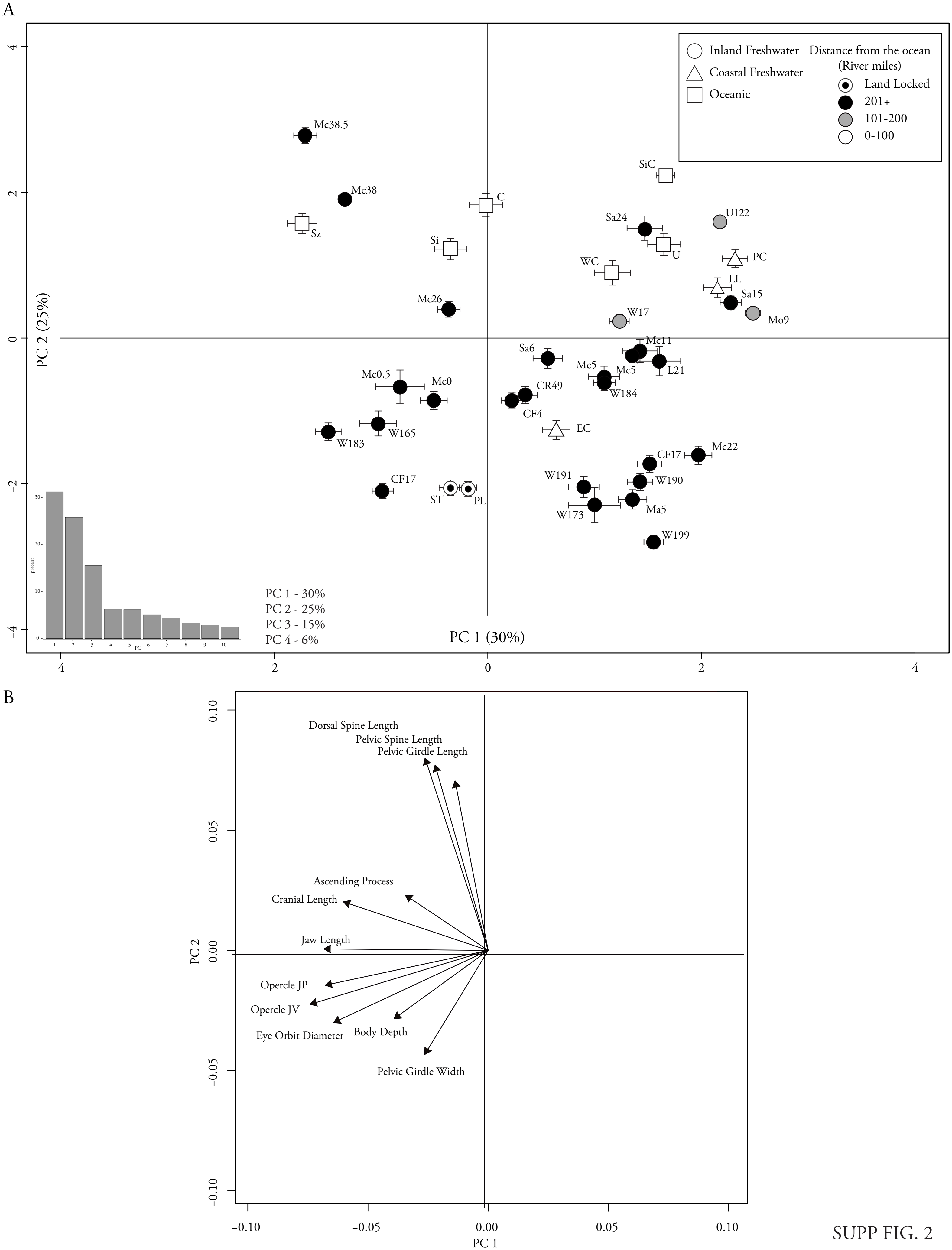

### Supplemental Fig. 3

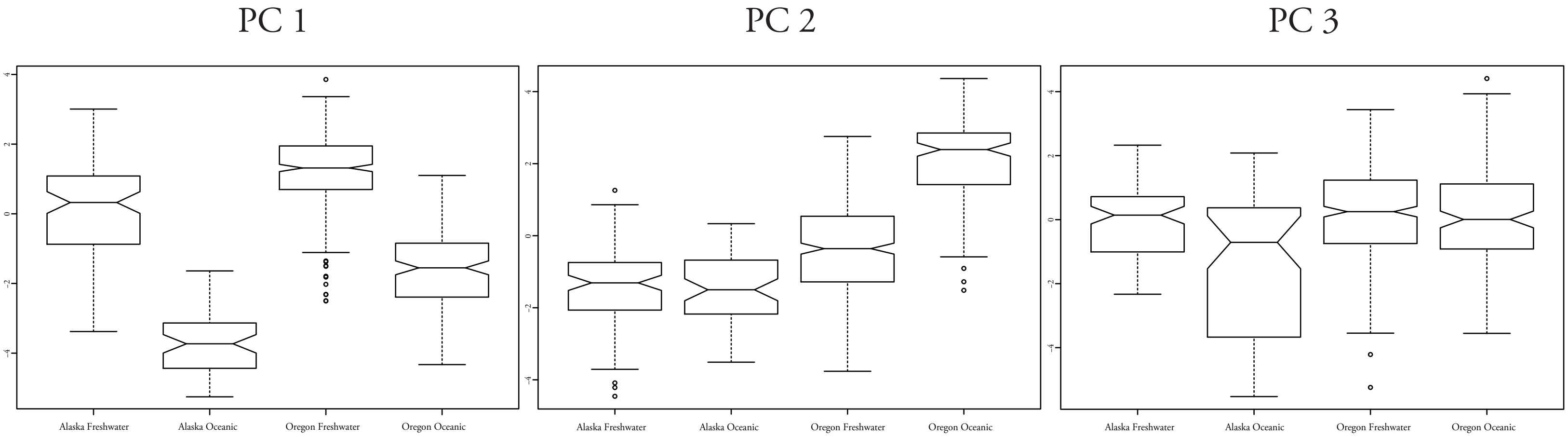

### Supplemental Fig. 4

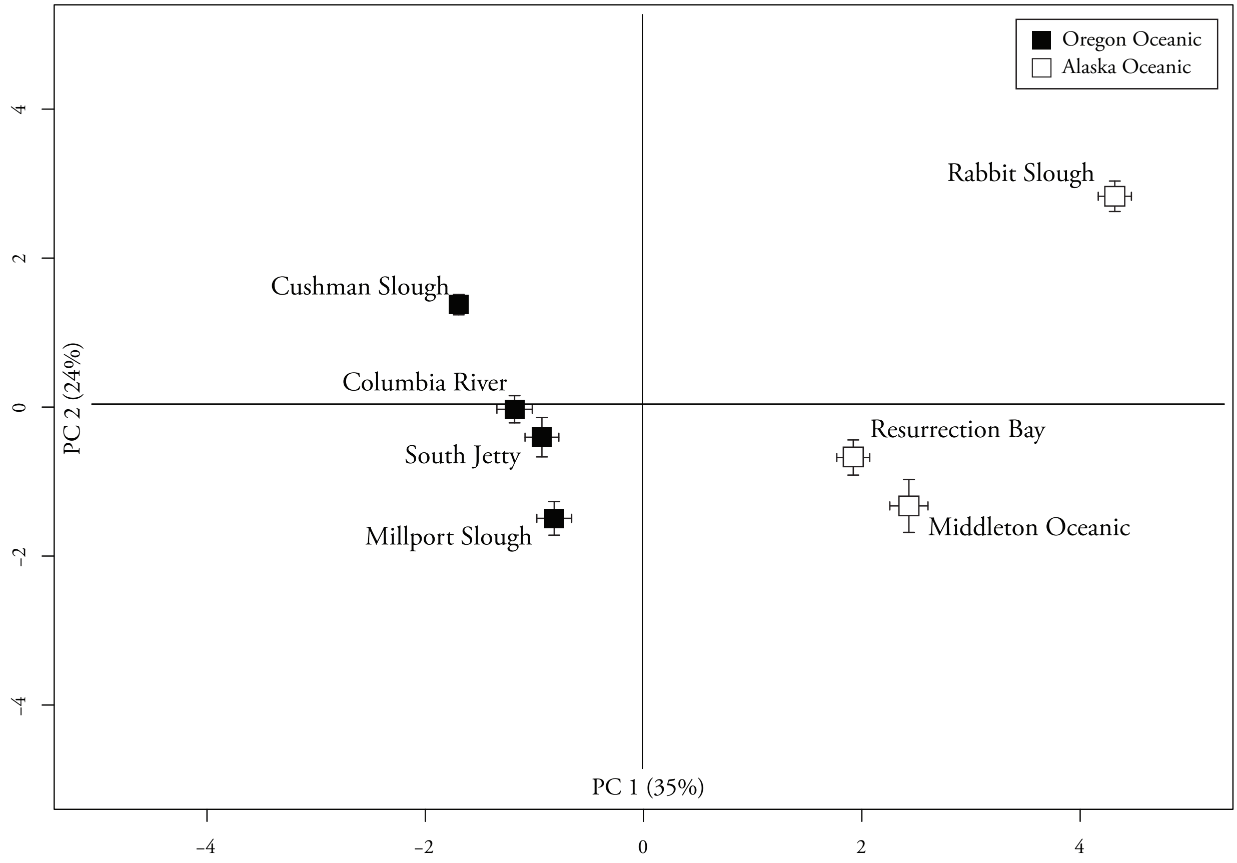

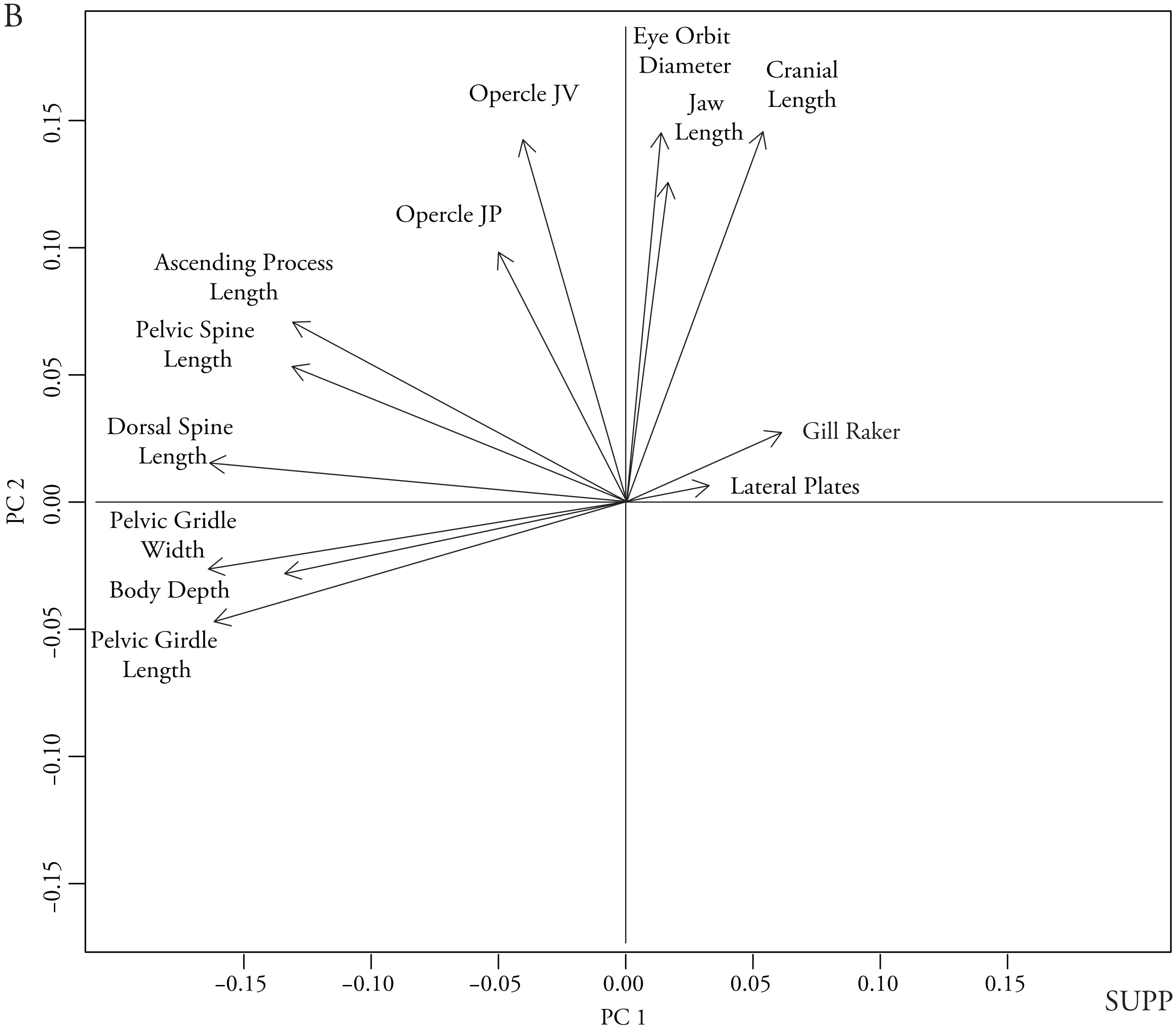
